# Supra-physiological levels of gibberellins/DELLAs modify the root cell size/number and the root architecture in root tips of *A. thaliana* seedlings. Connections to the root hair patterning and abundance

**DOI:** 10.1101/2021.07.25.453699

**Authors:** Iva McCarthy-Suárez

## Abstract

A previous study (McCarthy-Suárez, 2021) showed that growing *A. thaliana* seedlings for 5 days under excessive levels of gibberellins (GAs)/DELLAs altered the arrangement, shape and frequency of root hairs in root tips. Because no changes in the distribution or number of root hairs occurred when the gai-1 (gibberellin-insensitive-1) DELLA was over-expressed at the root epidermis, it was concluded that the GAs/DELLAs might regulate the root hair patterning and abundance in *A. thaliana* seedlings by acting from the root sub-epidermal tissues. In the present study, microscopy analyses showed that excessive levels of GAs/DELLAs also modified the size and number of root tip cells in *A. thaliana* seedlings. While excessive DELLAs shortened and widened the root epidermal, cortical, endodermal and pericycle cells, excessive GAs, excepting the epidermal cells, generally narrowed them. However, no changes of root cell size occurred when gai-1 was over-expressed at the root epidermis. In addition, high levels of DELLAs often induced extra cells at the root epidermis, cortex, endodermis and pericycle, whereas high levels of GAs sometimes induced extra cells at the root cortex and pericycle. On the other hand, excessive levels of DELLAs enhanced the outgrowth of lateral roots in root tips, unlike excessive levels of GAs. Thus, the results of this study suggest that supra-physiological levels of GAs/DELLAs might modify the size/number of root tip cells by acting from the root sub-epidermal tissues. This, in turn, might impact on the patterning and abundance of root hairs and on the root architecture.

## 1. INTRODUCTION

A previous study showed that supra-physiological levels of GAs/DELLAs altered the patterning, the morphology and the abundance of root hairs in *A. thaliana* seedlings, and that they did it by possibly acting from the sub-epidermal tissues of the root (McCarthy-Suárez, 2021). In fact, the GAs/DELLAs have a role in the production of trichomes (leaf hairs) in *A. thaliana* (Chien and Sussex, 1996; Traw and Bergelson, 2003) and participate in the organization of the cytoskeleton of microtubules (MT) (Locascio *et al*., 2013), which is essential for trichome and root hair growth, for establishing root cell identity and shape, and for plant cell expansion and division (Bao *et al*., 2001).

Because auxin, ethylene, abscisic acid, nitric oxide, brassinosteroids, cytokinins and strigolactones regulate the root hair patterning (Cao *et al*., 1999; Van Hengel *et al*., 2004; Lombardo *et al*., 2006; Kappusamy *et al*., 2009; Niu *et al*., 2011), and because changes in the levels of these phytohormones are correlated to alterations in the root cell size and number and the root architecture in response to nutritional stresses, such as low availability of P, B or Fe in the soil (e.g. swelling of root cortex cells, induction of extra root cortex cells and production of lateral roots (LR)) (Schmidt *et al*., 2000; Yang *et al*., 2007; Martín-Rejano *et al*., 2011), this study wanted to determine whether supra-physiological levels of GAs/DELLAs might also have an effect on the size/number of root cells, and on the production of LR, in root tips of *A. thaliana* seedlings. To this aim, the size and/or number of root tip cells were examined in Col (0) seedlings grown for 5 days under excessive levels of GAs/DELLAs, as well as in GAs (*QD, 5X*, GID*1b*-ox) or DELLAs (*gai*-1, *HSp::gai-1, pGAI:gai-1:GR, SCR::gai-1:GR*)-overproducing mutants. Moreover, the size of root tip cells was examined in 5-day-old mutant seedlings resulting from expressing the *gai-1* (GA-insensitive) DELLA allele in different tissues of the root (UAS (GAL4-UPSTREAM ACTIVATION SEQUENCE) expression directed system lines; Dr. Jim Haselhoffs laboratory). On the other hand, the presence of LR was also analysed in root tips of *A. thaliana* seedlings grown under (or harbouring) excessive levels of GAs/DELLAs.

Results of this study suggested that the GAs/DELLAs might have a role in regulating the size, number and organization of root cells, as well as the root architecture, in root tips of *A. thaliana* seedlings.

## 2. MATERIALS & METHODS

### 2.1. Plant Material and Growth Conditions

*Arabidopsis thaliana* Col (0) seeds were sterilized (70 % Ethanol (v/v) and 0.01 % Triton X-100 (v/v)), sown on half-strength MS medium plates (0.8 % (w/v) agar and 1 % (w/v) sucrose), stratified for 3-4 days (4°C, darkness), germinated, and grown vertically (22 °C; 5-7 days) under continuous white light (Percival growth chamber E-30B) (http://www.percival-scientific.com) as described by Lee and Schiefelbein (1999).

### 2.2. Hormone and Chemical treatments

Stock solutions of paclobutrazol (PAC, 10 mM in acetone 100 % (v/v)), GA_4_ (1 mM in 100 % ethanol (v/v)) or GA_3_ (50 mM in 100% ethanol (v/v)) were conveniently diluted and added to MS agar medium or water (in the case of liquid incubation experiments) to obtain a final concentration of 0.5 μM PAC, 1 μM GA_4_, and 30 μM GA_3_.

### 2.3. Mutant Lines

In the previous study, the spatial gene expression of the root non-hair epidermal cell fate marker GL2 in root tips of *A. thaliana* seedlings was studied by using the *GL2pro::GUS* mutant line as well as those derived from crossing lines harbouring constitutively excessive levels of GAs/DELLAs with the *GL2pro::GUS* line (L*er* x *GL2pro::GUS* background). In the present study, the effect of transient increases in the levels of expression of the *gai*-1 (GA-insensitive-1) DELLA alelle on the size and number of root tip cells in *A. thaliana* seedlings was examined by using the *gai-1* mutant lines of heat-shock inducible *HSp::gai-1* (which over-expresses gai-1 upon heat shock) and dexametasone (DEXA)-inducible *pGAI::gai-1:GR* and *SCR::gai-1:GR* (with glucocorticoid-binding domain). The *HSp::gai-1* mutant seedlings were grown at 37°C for 4 h (heat-shock) and then at 22°C for 2 h (recovery period), whereas the *pGAI::gai-1:GR* and *SCR::gai-1:GR* mutant seedlings were incubated in 0.2, 1.2 or 10 μM DEXA for a minimum of 6h. Root cell size and number was also studied in mutants with excessive levels of GAs/DELLAs (*gai*-1, *QD (quadruple DELLA mutant), 5X (quintuple DELLA mutant)* and GID1b-*ox* (which over-expresses the GA receptor GID1b (GIBBERELLIN INSENSITIVE DWARF1)), in mutants over-expressing gai-1 in different tissues of the root [*ML1::gai-1* (epidermis) and UAS expression directed system (GAL4-UPSTREAM ACTIVATION SEQUENCE) lines): *UAS::gai-1* x C24 (control, background); *UAS::gai-1* x J0951 (epidermis of the meristematic zone (MZ)); *UAS::gai-1* x J2812 (MZ epidermis and cortex); *UAS::gai-1* x N9142 (cortex of elongation zone (EZ)); *UAS::gai-1* x M0018 (MZ cortex and endodermis); *UAS::gai-1* x J0571 (MZ cortex and endodermis); *UAS::gai-1* x Q2393 (all tissues but the endodermis); *UAS::gai-1* x Q2500 (MZ endodermis/pericycle); *UAS::gai-1* x J0121 (EZ pericycle); *UAS::gai-1* x J0631 (all tissues of the EZ); *UAS::gai-1* x J3281 (vessels)], and in the *35S::CPC x GL2pro::GUS* and *scm* (scrambled) x *GL2pro::GUS* mutants.

### 2.4. GUS activity assay

GUS (β-glucuronidase) staining of the *GL2pro::GUS* reporter line was performed as described by Frigerio *et al*., (2006), but using 8mM instead of 2 mM potassium ferro/ferricyanide and incubating the seedlings (15 min to 2h) in the reaction mixture at 4 °C instead of 37 °C.

### 2.5. Microscopy

Cell organization at the root tip was studied on ultra-thin cross sections of plastic resin-embedded roots as described by Dr. Schiefelbein Lab Protocols (http://www.mcdb.lsa.umich.edu/labs/schiefel/protocols.html). Seedlings were included in 1% agarose in 0.1M sodium phosphate buffer, pH 6.8, and stained for GUS activity. Root-containing blocks were then cut, fixed with 4% para-formaldehyde in PBS, dehydrated in ethanol series (15%, 30%, 50%, 75%, 95% and 100%, 1h each), kept in 100% ethanol overnight, incubated in Technovit ^®^ 7100 infiltration solution for 2 days, inserted in gelatin capsules, and embedded for 9 days in Technovit ^®^ 7100 plastic resin (Heraeus Kultzer, Germany). Ultramicrotome (Ultracut E, Reichert Jung, Germany) cross sections of resin-embedded roots were then stained with 0.06% (w/v) toluidine blue and observed under a Nikon Eclipse E600 microscope. The GFP expression in Haselhoff crossed lines was visualized by using a Leica Confocal Microscope (Excitation: 488 nm; Detection: 500-530nm band-path filter for GFP).

## 3. RESULTS

### 3.1. Excessive levels of GAs/DELLAs modified the size and number of root tip cells in seedlings of *A. thaliana*

Apart from altering the patterning, morphology and abundance of root hairs (McCarthy-Suárez, 2021), high levels of GAs/DELLAs also modified the size and number of root tip cells in seedlings of *A. thaliana* (Figs. 1–3; Tables 1–3). While excessive levels of GAs, excepting the epidermal cells, usually narrowed the root cells, excessive levels of DELLAs frequently widened, shortened and twisted the root epidermal, cortical, endodermal and pericycle cells, resulting in wider roots (Figs. 1–3; Tables 1–3).

**Fig 1.**
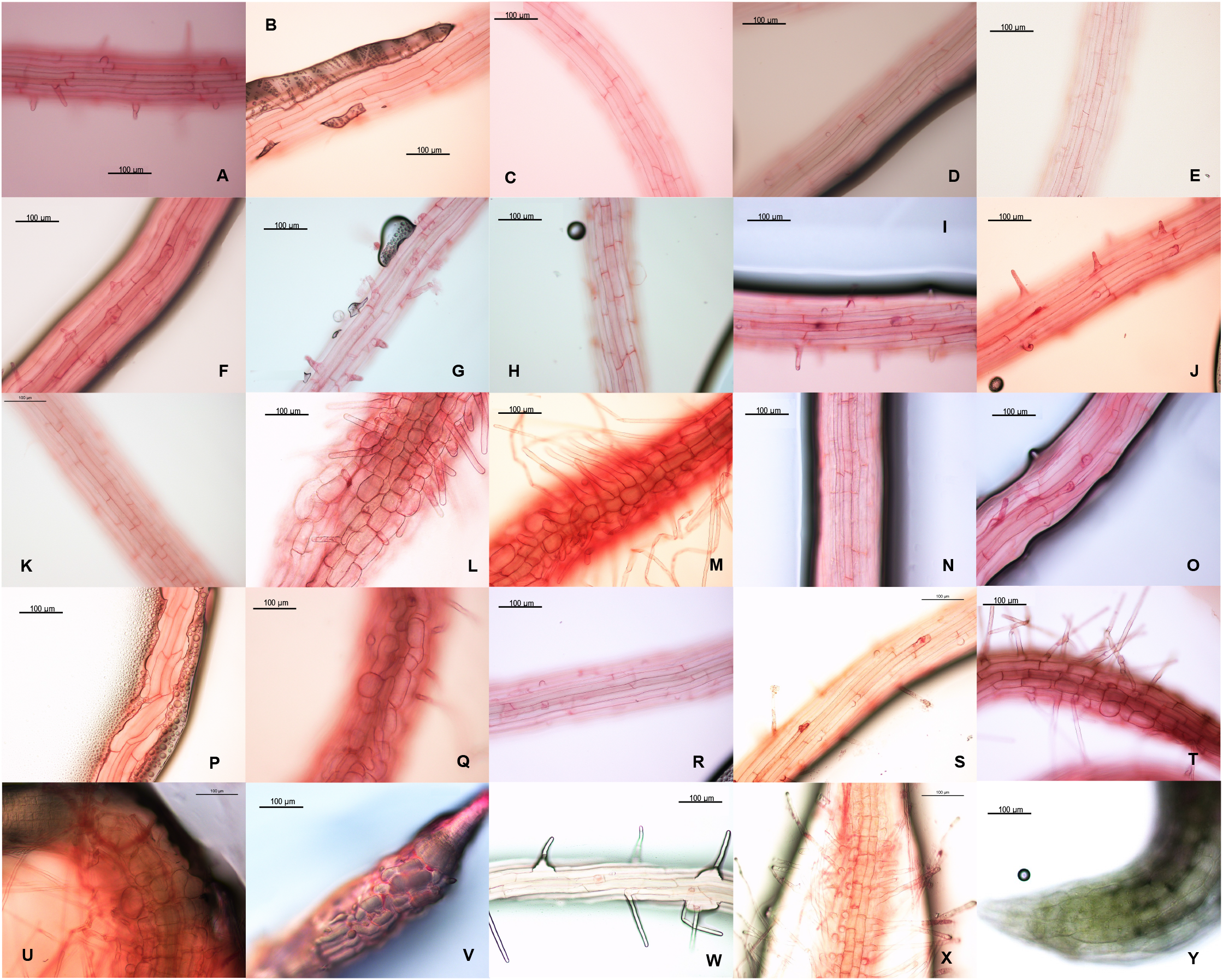
Excessive levels of GAs/DELLAs altered the size of root epidermal cells in root tips of 5-day-old A. thaliana seedlings. **A)** Col(0) (MS); **B)** Col(0) (0.5 μM PAC); **C)** Col(0) (30 μM GA_3_); **D)** Col (0) (1 μM GA_4_); **E)** L*er*; **F)** *gai-1*; **G)** *QD*; **H)** *5X*; **I)** GID1b-*ox* (MS); **J)** *HSp::gai-1* (22°C, 4h); **K)** *HSp::gai-1* (immediately after heat shock (37°C, 4h); **L)** *HSp::gai-1* (24h after heat-shock (37°C, 4h); **M)** *HSp::gai-1* (48h after heat-shock (37°C, 4h); **N)** *pGAI::gai-1:GR* (MS); **O)** *pGAI::gai-1:GR* (10 μM DEXA); **P)** *SCR::gai-1:GR* (MS, leaky line); **Q)** *SCR::gai-1:GR* (10 μM DEXA); **R)** *UAS::gai-1* x C24; **S)** *UAS::gai-1* x J0951; **T)** *UAS::gai-1* x J2812; **U)** *UAS::gai-1* x J0571; **V)** *UAS::gai-1* x Q2500; **W)** *UAS::gai-1* x J0121; **X)** *UAS::gai-1* x J0631; **Y)** *UAS::gai-1* x J3281. Magnification: 20X. Propidium iodide staining.

**Table 1.** Average length and width of root cortical cells in root tips of 5-day-old A. thaliana seedlings grown under (or harbouring) excessive levels of GAs/DELLAs. Analyses performed on electron micrographs of root cortex cells, 20X. (*) At 48h after a 4h-heat-shock experiment.

|  | Root cortical cell |  |  |  |
| --- | --- | --- | --- | --- |
|  | N° Cells analysed | Length (µm) | N° Cells analysed | Width (µm) |
| Col (0) (MS) | 25 | 171 ± 28 (100 %) | 66 | 31 ± 3 (100 %) |
| PAC (0.5 µM) | 23 | 153 ± 34 (89 %) | 60 | 43 ± 3 (142 %) |
| GA<sub>4</sub> (1 µM) | 26 | 209 ± 52 (122 %) | 26 | 27 ± 2 (87 %) |
| PAC (0.5 µM) + GA<sub>4</sub> (1 µM) | 24 | 173 ± 49 (101 %) | 28 | 19 ± 2 (61 %) |
| Ler | 28 | 188 ± 29 (100 %) | 18 | 30 ± 4 (100 %) |
| *ML1::gai-1* | 12 | 171 ± 33 (91 %) | 28 | 30 ± 3 (97 %) |
| *gai-1* | 71 | 138 ± 41 (73 %) | 72 | 37 ± 7 (123 %) |
| QD | 33 | 180 ± 34 (96 %) | 53 | 27 ± 3 (90 %) |
| 5X | 15 | 187 ± 34 (99 %) | 30 | 27 ± 4 (90 %) |
| *GID1b-ox* | 23 | 210 ± 54 (112%) | 34 | 24 ± 4 (80 %) |
| *Hsp::gai-1x GL2pro::GUS* (22°C) | 48 | 184 ± 39 (100 %) | 47 | 27 ± 4 (100 %) |
| *Hsp::gai-1 x GL2pro::GUS* (37°C) | 38 | 158 ± 52 (86 %) | 33 | 31 ± 4 (115 %) |
| *Hsp::gai-1 x GL2pro::GUS* (22°C) 48h* | 23 | 195 ± 43 (100 %) | 34 | 29 ± 3 (100%) |
| *Hsp::gai-1 x GL2pro::GUS* (37°C) 48h* | 36 | 61 ± 19 (31 %) | 44 | 46 ± 9 (159 %) |
| *pGAI::gai-1:GR* (MS) | 20 | 180 ± 56 (100 %) | 20 | 34 ± 4 (100 %) |
| *pGAI::gai-1:GR* (10 µM DEXA) | 29 | 129 ± 31 (72 %) | 29 | 46 ± 5 (135 %) |
| *SCR::gai-1:GR* (MS) | 77 | 140 ± 51 (100 %) | 77 | 34 ± 5 (100%) |
| *SCR::gai-1:GR* (10 µM DEXA) | 64 | 70 ± 32 (50 %) | 63 | 53 ± 10 (156 %) |
| *UAS::gai-1 x C24* (control) | 46 | 154 ± 49 (100 %) | 52 | 28 ± 4 (100 %) |
| *UAS::gai-1 x J0951* (epidermis) | 54 | 155 ± 42 (101%) | 72 | 31 ± 4 (111 %) |
| *UAS::gai-1 x J2812* (epi + cortex) | 79 | 116 ± 48 (75 %) | 91 | 35 ± 6 (125 %) |
| *UAS::gai-1 x J0571* (cortex + endo) | 74 | 71 ± 18 (46 %) | 74 | 48 ± 8 (171 %) |
| *UAS::gai-1 x M0018* (cortex + endo) | 49 | 89 ± 31 (58 %) | 46 | 46 ± 9 (164 %) |
| *UAS::gai-1 x Q2500* (MZ endo/pericycle) | 79 | 51 ± 13 (33 %) | 85 | 41 ± 8 (146 %) |
| *UAS::gai-1 x Q2393* (all but endo) | 23 | 174 ± 59 (113 %) | 32 | 32 ± 5 (114 %) |
| *UAS::gai-1 x J0631* (elong. tissues) | 73 | 46 ± 8 (30 %) | 67 | 37 ± 6 (132 %) |
| *UAS::gai-1 x J0121* (EZ pericycle) | 16 | 177 ± 45 (115 %) | 19 | 27 ± 3 (96 %) |

**Table 2.**
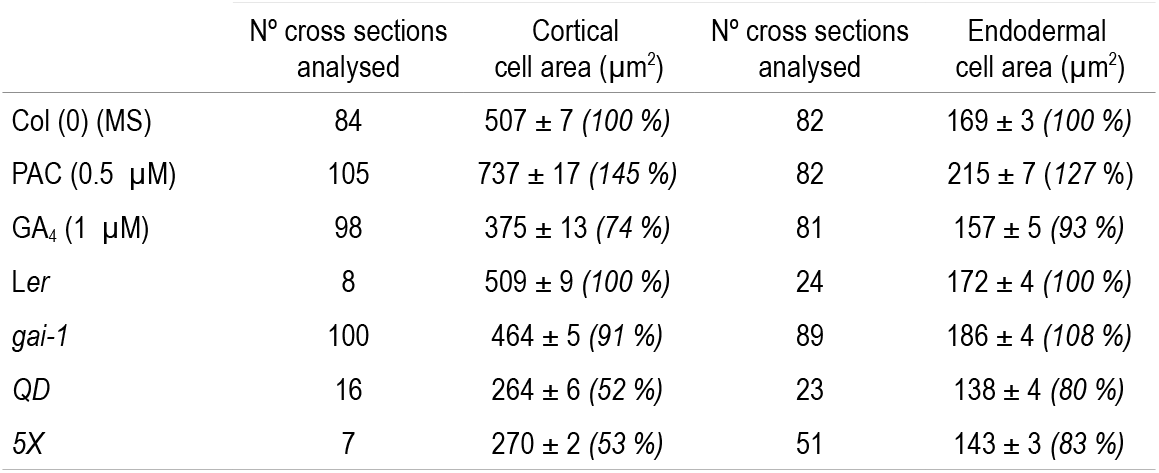
Cortical and endodermal cell area in cross sections of root tips of 5-day-old A. thaliana seedlings grown under (or harbouring) excessive levels of GAs/DELLAs. Measurements of transversal cell length and width were performed on electron micrographs of cross sections of resin-embedded roots (40X). Area was calculated by multiplying cell length (μm) by cell width (μm).

**Table 3.** Average width of the root MZ in root tips of 5-day-old A. thaliana seedlings grown under (or harbouring) excessive levels of GAs/DELLAs. Measurements of root width were performed on electron micrographs of whole root tips (10X, longitudinal view) or of cross sections of resin-embedded roots (40X).

| | Root width ($\mu\text{m}$) | | | |
| --- | --- | --- | --- | --- |
|  | N° roots analysed | Whole root tips | N° cross sections analysed | Root cross sections |
| Col (0) (MS) | 12 | 115 ± 4 (100 %) | 84 | 144 ± 10 (100 %) |
| PAC (0.5 $\mu\text{M}$) | 19 | 161 ± 12 (140 %) | 107 | 164 ± 14 (114 %) |
| GA<sub>4</sub> (1 $\mu\text{M}$) | 19 | 99 ± 12 (86 %) | 96 | 128 ± 18 (89 %) |
| Ler | 10 | 122 ± 5 (100 %) | 8 | 145 ± 4 (100 %) |
| *gai-1* | 18 | 125 ± 18 (102 %) | 100 | 148 ± 13 (102 %) |
| QD | 16 | 107 ± 18 (88 %) | 41 | 129 ± 7 (89 %) |
| 5X | 5 | 111 ± 0 (91 %) | 50 | 132 ± 8 (91 %) |

Moreover, in the *HSp::gai-1* and *SCR::gai-1:GR* mutants, the widening and shortening of the root epidermal, cortical, endodermal and pericycle cells observed at 24h after heat-shock (37°C, 4h) or after growth in DEXA (10 μM), respectively (Fig. 3), was accompanied by an alteration in the spatial expression of *GL2* and in the distribution of root hairs, as previously reported (McCarthy-Suárez, 2021). Similar changes were observed when gai-1 was over-expressed at the subepidermal tissues of the root (Figs. 1, 2 and 4; Table 1). However, when gai-1 was over-expressed at the root epidermis, no apparent changes occurred in the size of the root epidermal, cortical, endodermal or pericyclic cells (Figs. 1, 2 and 4; Table 1), in the patterning of GL2 gene expression, or the in root hair distribution (McCarthy-Suárez, 2021), what suggested that the changes in root cell size that took place when gai-1 was over-expressed at the sub-epidermal tissues of the root might have been connected to the alterations in the root hair patterning induced by excessive levels of GAs/DELLAs.

**Fig 2.**
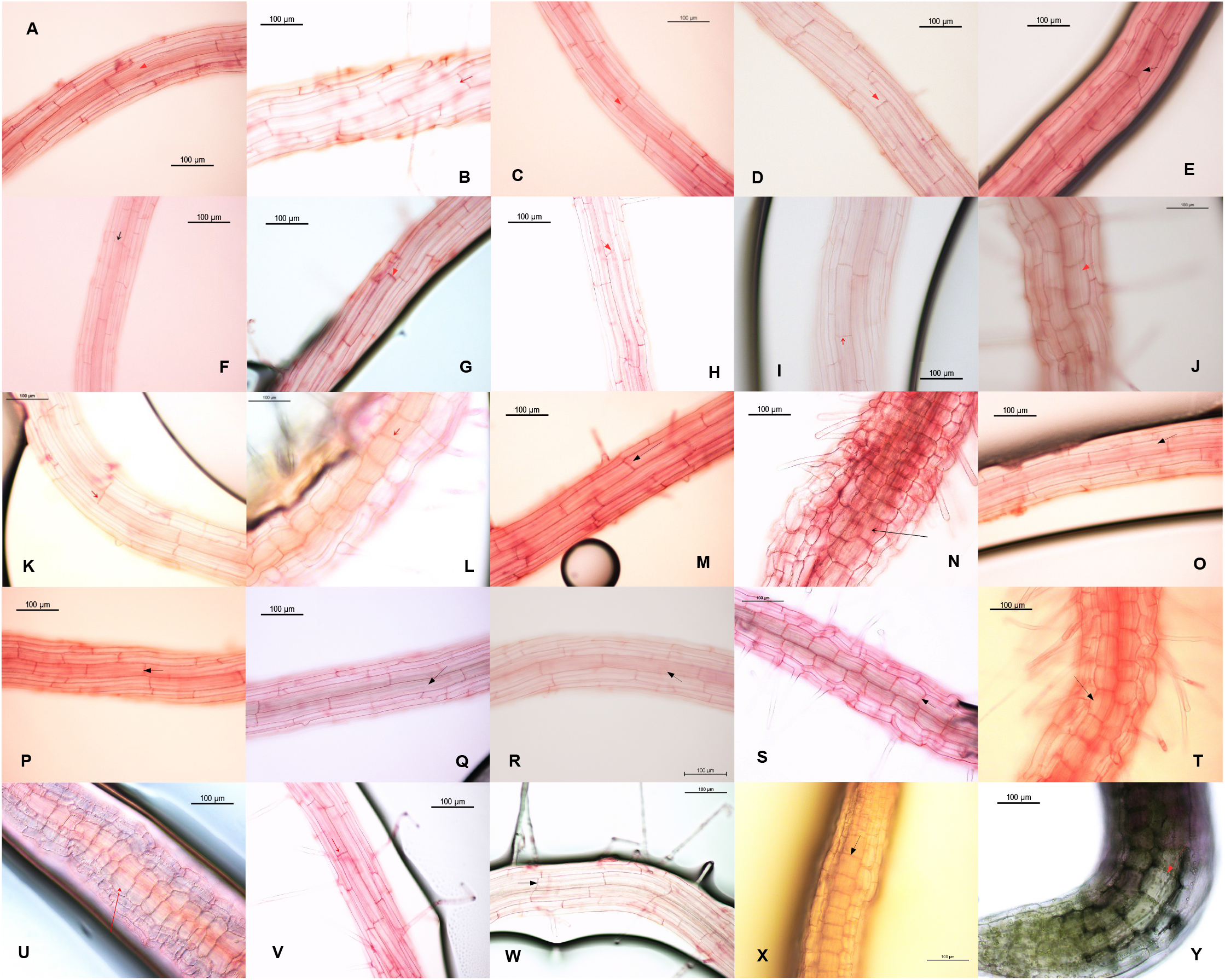
Excessive levels of GAs/DELLAs altered the size of root cortical cells in root tips of 5-day-old A. thaliana seedlings. **A)** Col(0) (MS); **B)** Col(0) (0.5 μM PAC); **C)** Col(0) (1 μM GA_4_); **D)** L*er*; **E)** *gai-1;* **F)** *QD*; **G)** *5X*; **H)** GID*1b-ox;* **I)** *pGAI::gai-1:GR* (3d in MS); **J)** *pGAI::gai-1:GR* (3d in 10 μM DEXA); **K)** *SCR::gai-1:GR* (3d in MS); **L)** *SCR::gai-1:GR* (3d in 10 μM DEXA); **M)** *HSp::gai-1* x *GL2pro::GUS* (24h after 22°C for 4h); **N)** *HSp::gai-1* x *GL2pro::GUS* (24h after heat-shock (37°C for 4h)); **O)** *GL2pro::GUS* (heat-shock control) (24h after 22°C for 4h); **P)** *GL2pro::GUS* (heat-shock control) (24h after heat shock (37°C for 4h)); **Q)** *UAS::gai-1* x C24; **R)** *ML1::gai-1;* **S)** *UAS::gai-1* x J2812; **T)** *UAS::gai-1* x M0018; **U)** *UAS::gai-1* x Q2500; **V)** *UAS::gai-1* x N9142; **W)** *UAS::gai-1* x J0121; **X)** *UAS::gai-1* x 0631; **Y)** *UAS::gai-1* x J3281. Magnification: 20X. Propidium iodide staining.

Growth of *A. thaliana* seedlings for 5 days under excessive levels of GAs, in contrast, caused the narrowing of the root cortical cells, an effect that was corroborated in the *QD, 5X* and GID1b *ox* mutants (Fig. 2; Table 1). Nevertheless, excessive levels of GAs also seemed to slightly increase the relative width of the root epidermal cells (Figs. 1, 5 and 6). Frequently, under GA_3_ (30 μM) or GA_4_ (1 μM), changes of cell fate at the root epidermis coincided with changes in the width of the epidermal cells and/or with changes in the size of the root cortical and endodermal cells (Fig. 3; Tables 1 and 2). In fact, estimations of tissue depth in root tips of *A. thaliana* seedlings uncovered the swelling of the root epidermal, cortical, endodermal and pericyclic cells under high levels of DELLAs (PAC) and their slight thinning under high levels of GAs (*5X* mutant) (Table 4). Moreover, growth of seedlings of the *scm* x *GL2pro::GUS* mutant in PAC (0.5 μM) for 5d caused the radial swelling of the epidermal, cortical and endodermal and pericyclic cells of the root tip (Fig. 6). On the other hand, in seedlings of the *35S::CPC* x *GL2pro::GUS* mutant, changes of cell fate at the root epidermis were accompanied by changes in the width of the underlying root epidermal and cortical cells (Fig. 3).

**Fig 3.**
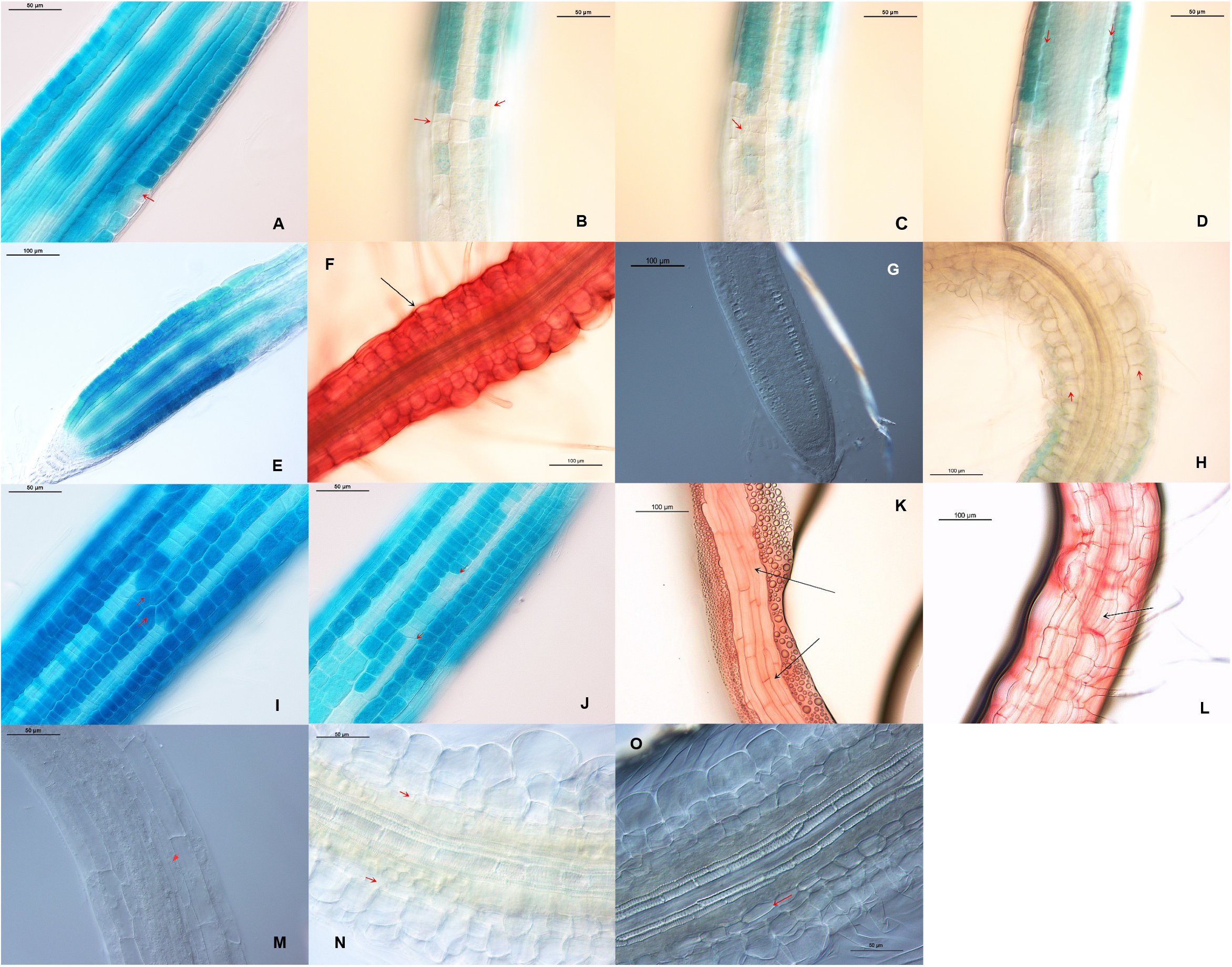
Excessive levels of GAs/DELLAs induced cell size changes and T-clones at the epidermis, cortex and endodermis of root tips of 5-day-old A. thaliana seedlings. **A)** *GL2pro::GUS* (30 μM GA_3_) (all layers): expansion of an epidermal cell and narrowing of a cortical cell coincide with a change of epidermal cell fate; **B)** *35S::CPC* x *GL2pro::GUS* (epidermis): A change in epidermal cell size coincides with a change in epidermal cell fate; **C)** *35S::CPC* x *GL2pro::GUS* (cortex): A change in cortical cell width coincides with a change in epidermal cell fate; **D)** *35S::CPC* x *GL2pro::GUS* (all layers): epidermis and cortex vary in cell size; **E)** *HSp::gai-1* x *GL2pro::GUS* (48h after 22°C for 4h) (all layers), 20X; **F)** *HSp::gai-1* x *GL2pro::GUS* (48h after heat-shock (37°C, 4h) (all layers): swelling of the root epidermal, cortical, endodermal and pericyclic cells, 20X; **G)** *SCR::gai-1:GR* (MS) (all layers), 20X; **H)** *SCR::gai-1:GR* x *GL2pro::GUS* (24h in 10 μM DEXA) (all layers): swelling of the root cortex, endodermis and pericycle, 20X; **I)** Epidermal T-clones in PAC (0.5 μM), 40X; **J)** Epidermal T-clones in GA_3_ (30 μM), 40X; **K)** *SCR::gai-1:GR* (0.2 μM DEXA): epidermal T-clones, 20X; **L)** *SCR::gai-1:GR* (1.2 μM DEXA): periclinal cell division at the cortex, 20X; **M)** *SCR::gai-1:GR* (MS) (all layers), 40X: arrow on endodermis; **N)** *SCR::gai-1:GR* (10 μM DEXA) (all layers): periclinal cell divisions at the endodermis, 40X; **O)** *SCR::gai-1:GR* (10 μM DEXA) (all layers): periclinal cell divisions at the endodermis, 40X; Propidium iodide or GUS staining.

**Table 4.** Estimated tissue depth in roots tips of 5-day-old A. thaliana seedlings grown under (or harbouring) excessive levels of GAs/DELLAs. Measurements of tissue diameter (μm) were performed on electron micrographs of cross sections of resin-embedded roots (40X; MZ to early EZ). Estimated epidermal depth = [Root diameter (data from Table 3) – (Cortex-Endodermis-Pericycle-Vessels diameter)]/2. Estimated cortical depth = [(Cortex-Endodermis-Pericycle-Vessels diameter) – (Endodermis-Pericycle-Vessels diameter)]/2. Estimated endodermal depth = [(Endodermis-Pericycle-Vessels diameter) – (Pericycle-Vessels diameter)]/2. Estimated pericycle depth = [(Pericycle-Vessels diameter) – (Vessels diameter)]/2. Endo: Endodermis; Peri: Pericycle. Number of cross sections analysed: Control (27-29), PAC (46-49), GA_4_ (74-78), L*er* (21-22), *gai*-1 (36), *QD* (34-36), *5X*(47-48).

| | Col (0)<br>(MS) | PAC<br>(0.5 $\mu\text{M}$) | GA<sub>4</sub><br>(1 $\mu\text{M}$) | *Ler* | *gai-1* | *QD* | *5X* |
| --- | --- | --- | --- | --- | --- | --- | --- |
| Cortex-Endo-Peri-Vessels diameter | 95 ± 6<br>(100 %) | 110 ± 11<br>(116 %) | 83 ± 12<br>(87 %) | 90 ± 7<br>(100 %) | 91 ± 5<br>(101 %) | 81 ± 5<br>(90 %) | 82 ± 4<br>(91 %) |
| Endo-Peri-Vessels diameter | 59 ± 6<br>(100 %) | 70 ± 7<br>(119 %) | 58 ± 7<br>(98 %) | 60 ± 4<br>(100 %) | 59 ± 3<br>(98 %) | 59 ± 6<br>(98 %) | 55 ± 2<br>(92 %) |
| Peri-Vessels diameter | 45 ± 5<br>(100 %) | 54 ± 5<br>(120 %) | 45 ± 6<br>(100 %) | 45 ± 3<br>(100 %) | 44 ± 2<br>(98 %) | 45 ± 5<br>(100 %) | 40 ± 2<br>(89 %) |
| Vessels diameter | 35 ± 3<br>(100 %) | 40 ± 3<br>(114 %) | 32 ± 5<br>(91 %) | 33 ± 3<br>(100 %) | 34 ± 2<br>(103 %) | 33 ± 4<br>(100 %) | 30 ± 2<br>(91 %) |
| Estimated Epidermal depth | 25<br>(100 %) | 27<br>(108 %) | 22<br>(88 %) | 27<br>(100 %) | 28<br>(104 %) | 24<br>(89 %) | 25<br>(93 %) |
| Estimated Cortex depth | 18<br>(100 %) | 20<br>(111 %) | 13<br>(72 %) | 15<br>(100 %) | 16<br>(107 %) | 11<br>(73 %) | 13<br>(87 %) |
| Estimated Endodermis depth | 7<br>(100 %) | 8<br>(114 %) | 6<br>(86 %) | 8<br>(100 %) | 7<br>(88 %) | 7<br>(88 %) | 7<br>(88 %) |
| Estimated Pericycle depth | 5 (100 %) | 7 (140 %) | 6 (120 %) | 6 (100 %) | 5 (83 %) | 6 (100 %) | 5 (83 %) |

Interestingly, cell size changes in root tips under excessive levels of GAs/DELLAs were often accompanied by the presence of multinucleated cells at the epidermis, cortex, endodermis and pericycle of the root (Fig. 6).

Excessive levels of GAs/DELLAs also modified the radial cell organization in root tips of *A. thaliana* seedlings (Fig. 5). Treatments with PAC (5d, 7d) frequently increased the number of cells at the epidermis, cortex, endodermis and pericycle of the root (Figs. 5 and 6; Table 5) and induced anticlinal /diagonal cell divisions at the root epidermis (T-clones) (Fig. 3) as well as periclinal cell divisions at the root cortex and endodermis (middle cortex (MC)) (Fig. 6). Furthermore, growth of the *scm* x *GL2pro::GUS* mutant for 5 days in PAC induced the proliferation of the root epidermal, cortical and endodermal cells and the formation of a MC (Fig. 6). Treatments with excessive levels of GAs, in turn, sometimes increased the number of cortical and pericycle cells, but not of epidermal cells, in the radial dimension of the root, and induced epidermal T-clones and a MC (Figs. 3, 5 and 6; Table 5). Moreover, the number of root epidermal cells in the *5X* mutant decreased (Fig. 5; Table 5), what in part might have explained the lower abundance of root hairs per root radial section in this mutant (McCarthy-Suárez, 2021). In fact, in root cross sections, frequently only one epidermal cell was seen at the atrichoblast (non-hair) position under high levels of GAs, whereas up to four cells could be seen under high levels of DELLAs (Fig. 5), in tune with the reduced number of epidermal cells in the *5X* mutant and the increased number of epidermal cells under PAC (Table 5). Furthermore, given that root non-hair cells lay over just one cortical cell, then, the observed increase in the cortex cell width under high levels of DELLAs (Figs. 2, 4 and 5; Table 1) might have accounted for the higher percentage of epidermal cells at the atrichoblast position, as well as the lower percentage of epidermal cells at the trichoblast position, detected per radial section of the root (McCarthy-Suárez, 2021). Conversely, the decrease in the width of the cortex cells seen under high levels of GAs (*5X* mutant) (Fig. 2; Table 1) might have explained the lower percentage of epidermal cells at the atrichoblast position, and the higher percentage of epidermal cells at the trichoblast position, found per radial section of the root (McCarthy-Suárez, 2021). Nevertheless, considering that the average number of epidermal cells per root radial section increased under high levels of DELLAs (PAC, *gai*-1) and decreased under high levels of GAs (*5X* mutant) (McCarthy-Suárez, 2021), then, the predicted number of epidermal cells at the trichoblast position did not change under excessive levels of GAs/DELLAs in 5 day-old *A. thaliana* seedlings (McCarthy-Suárez, 2021).

**Fig 4.**
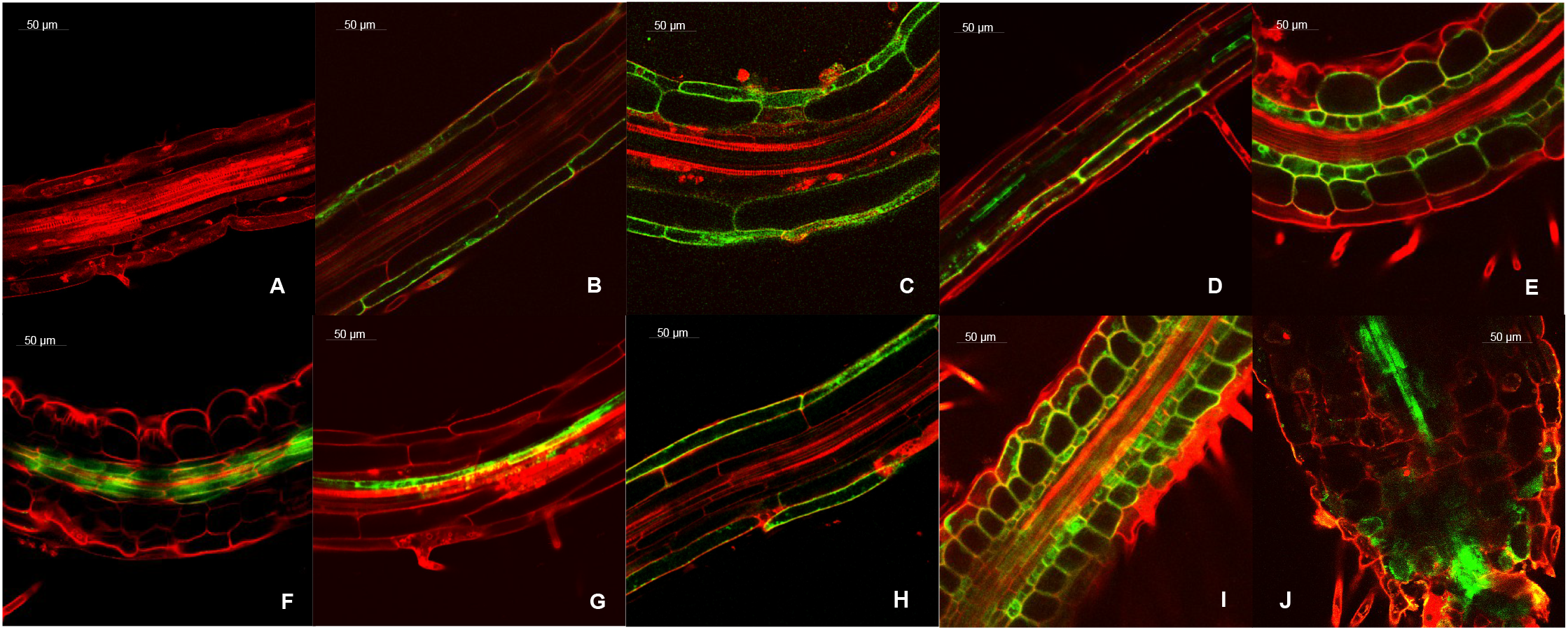
Over-expression of gai-1 in different tissues of the root modified the cell size in root tips of 5-day-old A. thaliana seedlings. **A)** *UAS::gai-1* x C24 (Background); **B)** *UAS::gai-1* x J0951 (epidermis of MZ); **C)** *UAS::gai-1* x J2812 (epidermis and cortex of the MZ); **D)** *UAS::gai-1* x N9142 (cortex of EZ); **E)** *UAS::gai-1* x J0571 (cortex and endodermis); **F)** *UAS::gai-1* x Q2500 (endodermis/pericycle of MZ); **G)** *UAS::gai-1* x J0121 (pericycle of EZ); **H)** *UAS::gai-1* x Q2393 (all tissues but the endodermis); **I)** *UAS::gai-1* x J0631 (elongating tissues); **J)** *UAS::gai-1* x J3281 (vessels). Magnification: 40X. Propidium iodide staining.

**Table 5.** Radial cell organization in roots of 5 or 7-day-old A. thaliana seedlings grown under (or harbouring) excessive levels of GAs/DELLA. Analyses performed on electron micrographs of cross sections of resin embedded-roots (40X).

|  | N° root cross sections analysed | Epidermis | Cortex | Endodermis | Pericycle |
| --- | --- | --- | --- | --- | --- |
| Control (5d) | 75 | 20–25 | 8 | 8–10 | 13–15 |
| PAC (5d) | 56 | 25–34 | 8–9 | 9–13 | 14–18 |
| GA<sub>4</sub> (5d) | 56 | 17–25 | 8–9 | 8–10 | 12–16 |
| Control (7d) | 14 | 24–27 | 8 | 8–10 | 14–15 |
| PAC (7 d) | 18 | 26–33 | 8–10 | 11–13 | 14–18 |
| GA<sub>4</sub> (7d) | 45 | 23–25 | 8–9 | 8–10 | 14–16 |
| Ler (5d) | 13 | 18–24 | 8 | 8 | 13–14 |
| *gai-1* (5d) | 57 | 19–25 | 8 | 8–9 | 12–14 |
| QD (5d) | 24 | 22–24 | 8–11 | 8–11 | 14–18 |
| 5X (5d) | 54 | 18–21 | 8–9 | 7–8 | 12–16 |

The root diameter at the MZ, in addition, increased by 40% under excessive levels of DELLAs (Table 3), in tune with the increased number of cells at the root epidermis, cortex, endodermis and pericycle (Fig. 5; Table 5), and the wider and/or deeper cells at the root cortex, endodermis and pericycle (Figs. 5 and 6; Tables 1 and 4). Conversely, under excessive levels of GAs, the root diameter decreased (Table 3), in accordance with the lower number of cells at the root epidermis (*5X* mutant) (Table 5), and the narrower and shallower cells at the root cortex and endodermis (Figs. 2, 5 and 6; Tables 1 and 4). Nevertheless, at the MZ-EZ transition zone, the root also seemed to swell under excessive levels of GAs, and there was variability of cell sizes (McCarthy-Suárez, 2021), maybe due to the swelling of the epidermal cells and to the deepening of the pericycle cells (Fig. 1; Table 4). In fact, GA_3_ treatments have been shown to increase the ratio of (xilem/whole root) area (Wang *et al*., 2015).

**Fig 5.**
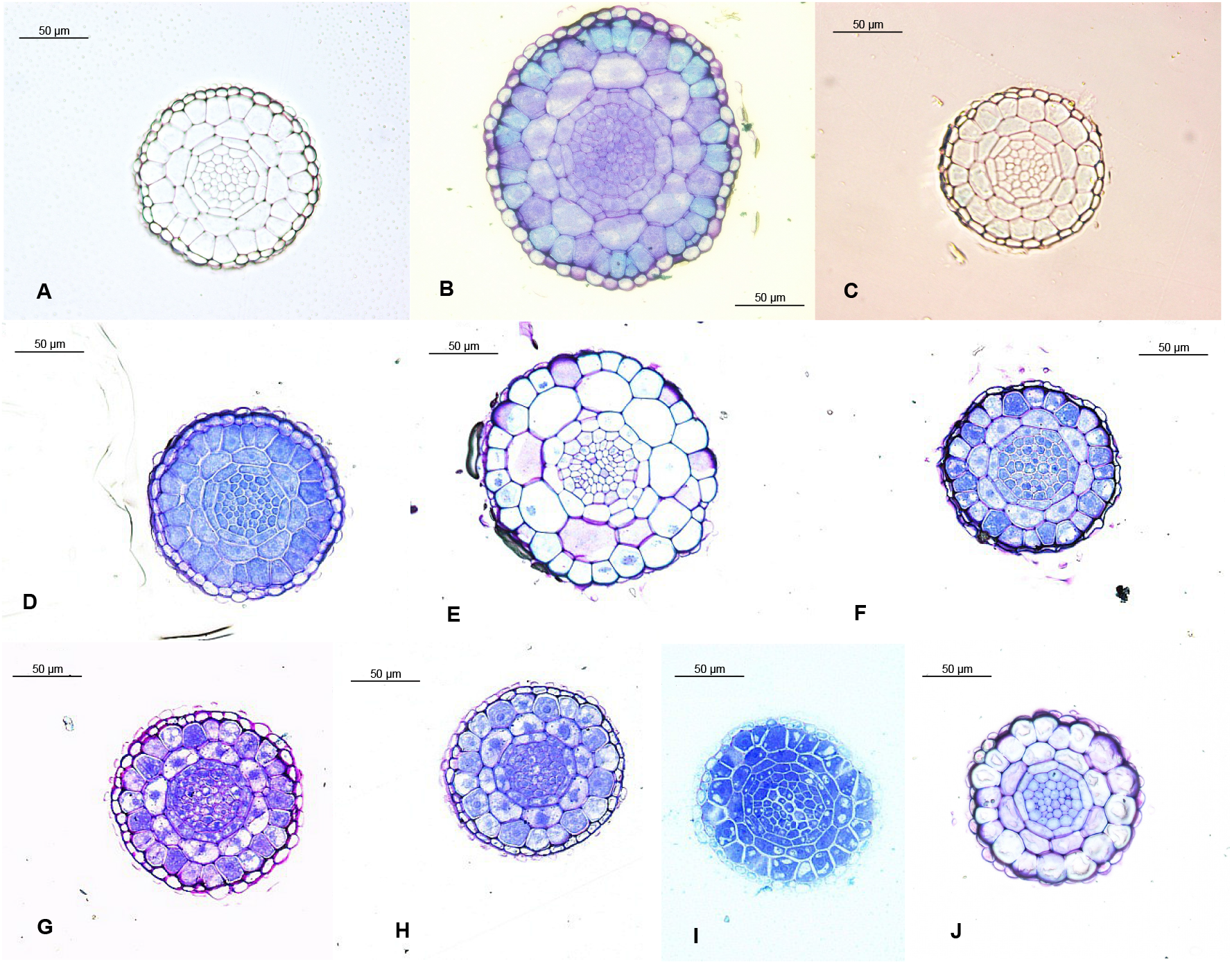
Radial cell organization in root tips of 5 or 7 day-old A. thaliana seedlings grown under (or harbouring) excessive levels of GAs/DELLAs. **A)** Col(0) (MS, 5d); **B)** Col(0) (0.5 μM PAC, 5d); **C)** Col(0) (1 μM GA_4_, 5d); **D)** Col(0) (MS, 7d); **E)** Col(0) (0.5 μM PAC, 7d); **F)** Col(0) (1 μM GA_4_, 7d); **G)** L*er* (5d); **H)** *gai*-1 (5d); **I)** *QD* (5d); **J)** *5X*(5d). Magnification: 40X. Toluidine blue staining.

**Fig 6.**
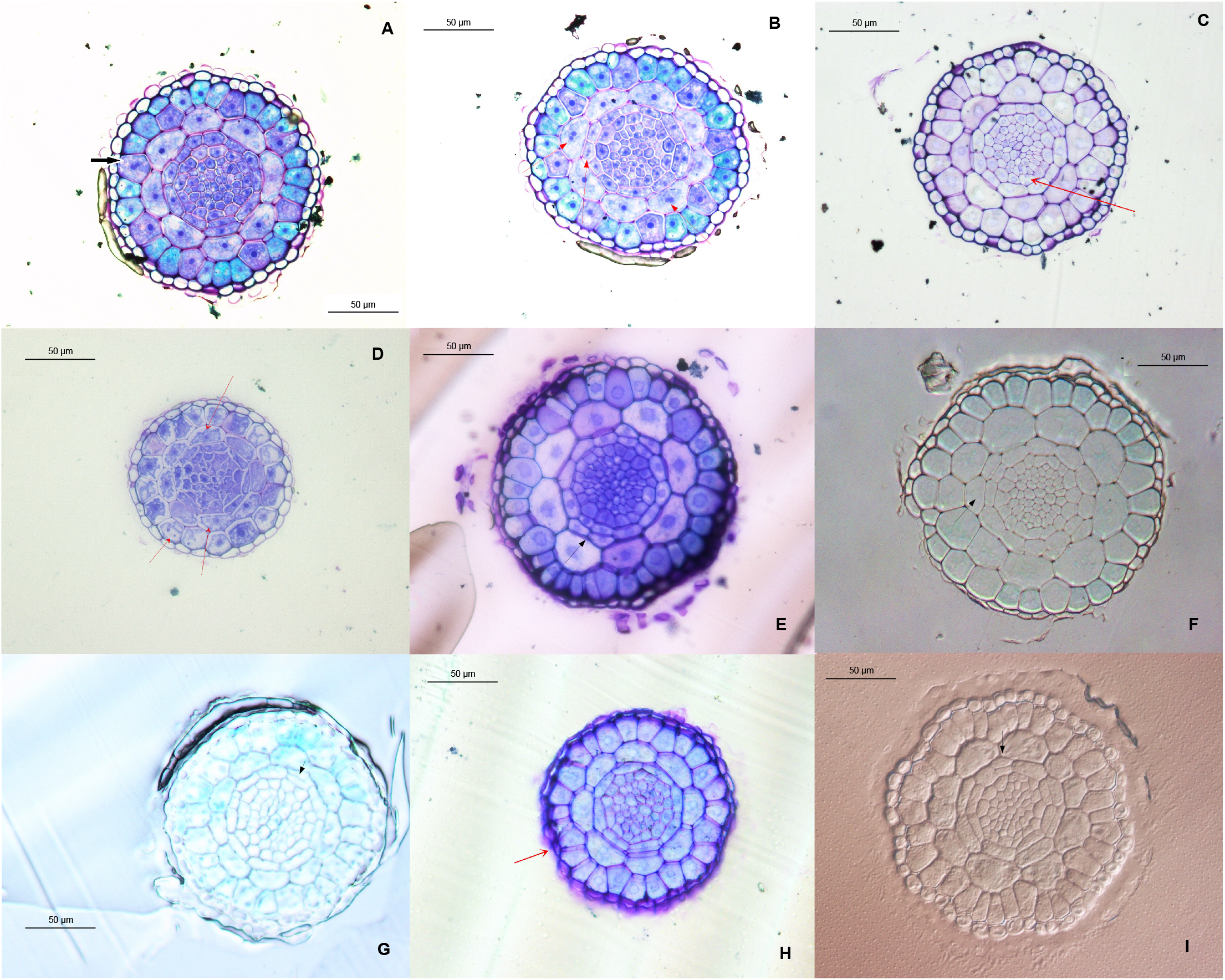
Excessive levels of GAs/DELLAs induced multinucleated cells, a middle cortex (MC) and extra cortical cells in root tips of A. thaliana seedlings. **A)** PAC (0.5 μM, 5d): Epidermal multinucleated cell; **B)** PAC (0.5 μM, 5d): Cortical and endodermal multinucleated cells; **C)** PAC (0.5 μM, 5d): Pericyclic multinucleated cell; **D)** GA_4_ (1 μM, 5d): Epidermal and cortical multinucleated cells; **E)** PAC (0.5 μM, 5d): MC; **F)** PAC (0.5 μM, 7d): MC (arrow) and 10 cortical cells; **G)** GA_4_ (1 μM, 5d): MC (arrow) and 9 cortical cells; **H)** *scm* x *GL2pro::GUS* (MS, 5d); **I)** *scm* x *GL2pro::GUS* (0.5 μM PAC, 5d): MC (arrow) and 9 cortical cells. Magnification: 40X. Toluidine blue staining.

### 3.2. Excessive levels of GAs/DELLAs modified the outgrowth of lateral roots in root tips of *A. thaliana* seedlings

Excessive levels of DELLAs also induced the outburst of LR near the root tip in *A. thaliana* seedlings, whereas excessive levels of GAs inhibited it (Fig. 7).

**Fig 7.**
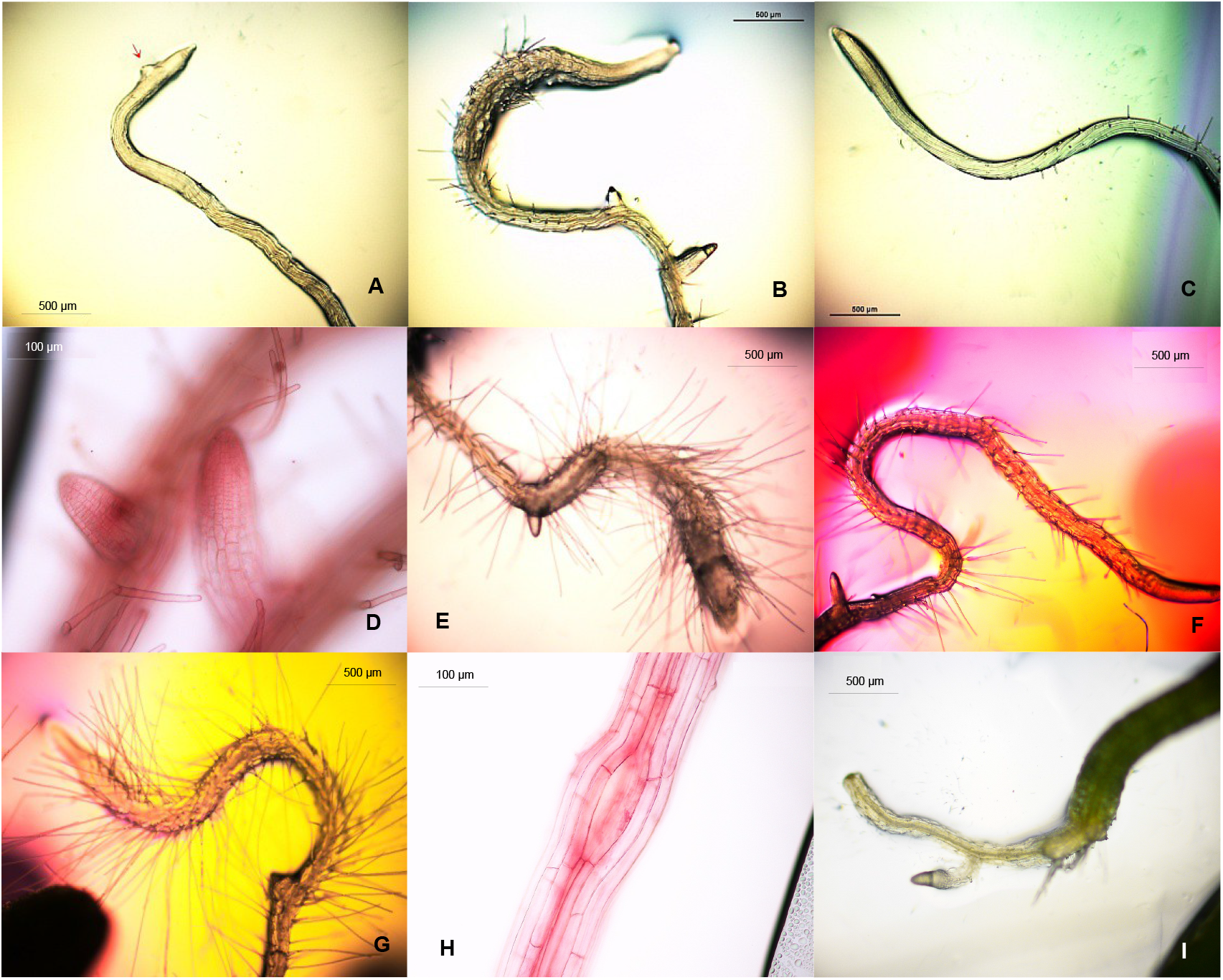
Over-expression of gai-1 induced an early outburst of lateral roots in root tips of A. thaliana seedlings. **A)** *SCR::gai-1:GR* (MS, 5d) (Leaky line), 4X; **B)** *SCR::gai-1:GR* (10 μM DEXA, 5d), 4X; **C)** *UAS::gai-1* x C24 (control, 5d), 4X; **D)** *UAS::gai-1* x J2812 (5d), 20X; **E)** *UAS::gai-1* x M0018 (5d), 4X; **F)** *UAS::gai-1* x J0571 (5d), 4X; **G)** *UAS::gai-1* x Q2500 (5d), 4X; **H)** *UAS::gai-1* x N9142 (5d), 20X; **I)** *UAS::gai-1* x J3281 (8d) (aborted primary root), 4X.

## 4. DISCUSSION

### 4.1. The GAs/DELLAs might regulate the size and number of root tip cells in seedlings of *A. thaliana*

#### 4.1.1. Root cell size. Connections to the root hair patterning and abundance

Apart from altering the patterning, morphology and abundance of root hairs (McCarthy-Suárez, 2021), results of the present study suggest that excessive levels of GAs/DELLAs also modified the size of root tip cells in seedlings of *A. thaliana*. While excessive levels of DELLAs frequently shortened and widened the epidermal, cortical, endodermal and pericyclic cells of the root, what resulted in wider root tips, excessive levels of GAs, with the exception of the epidermal cells, often narrowed them. Thus, because root hair cells are shorter than the root non-hair cells (Salazar-Henao *et al*., 2016), then, the inhibition of epidermal cell elongation that occurred when *gai-1* was over-expressed in tissues placed underneath the epidermis of the MZ (J2812 >> *gai-1*, J0571 >> *gai-1*, M0018 >> *gai-1*, Q2500 >> *gai-1* or Q2393 >> *gai-1* lines) or in all tissues of the EZ (J0631 >> *gai-1* line) might have contributed, in part, to the appearance of ectopic root hairs, and, therefore, to the higher density of root hairs observed near the root tip under excessive levels of DELLAs (McCarthy-Suárez, 2021). In fact, *Arabidopsis* increases root hair density by decreasing the length of root epidermal cells, as shown under P deficiency (Jiang *et al*., 2007; Péret *et al*., 2011; Salazar-Henao *et al*., 2016; Janes *et al*., 2018). Alternatively, given that GAs promote the elongation of root epidermal cells, accumulate at the endodermis of the root EZ, and affect the expansion of root EZ cells by destabilizing DELLAs and inducing expansin genes (Ubeda-Tomás *et al*., 2008, 2009; Gou *et al*., 2010; Bahin *et al*., 2011; Shani *et al*., 2013), then, the extra elongation of root epidermal cells which is known to occur under high levels of GAs (Band *et al*., 2012) might have contributed, in turn, to the lower density of root hairs that was observed at the root tip under this treatment (McCarthy-Suárez, 2021). In fact, in WT, cell expansion at the root EZ is strictly polar and is not accompanied by an increase in the root diameter (Bao *et al*., 2001).

Interestingly, epidermal patterning genes instruct epidermal cell size (Löfke *et al*., 2013). In turn, variations in the expression of HDA 19 (histidine deacetylase 19), which controls epidermal cell elongation, affect root cell elongation and, thus, root hair density (Chen *et al*., 2015; Salazar-Henao *et al*., 2016). In this study, however, neither the patterning or abundance of root hairs (McCarthy-Suárez, 2021), nor the epidermal cell length suffered alterations when *gai-1* was over-expressed at the root epidermis (Fig. 4), what suggests that the changes of epidermal cell size that were induced by excessive levels of GAs/DELLAs in roots of *A. thaliana* seedlings (Fig. 1) might have been orchestrated from the sub-epidermal tissues of the root. Furthermore, Wild *et al*. (2016) showed that expressing *gai-1* at the root epidermis did not affect the root length.

Results of this study also seem to point at a link between cell size and cell fate, because expressing *gai-1* at the EZ of the root (J0631 >> *gai-1)* caused the shortening and widening of the majority of the root cells (Fig. 4), along with a hairy phenotype similar to that of the *wer* mutant (McCarthy-Suárez, 2021). In fact, the DELLAs inhibit the elongation of the root EZ cells and of the primary root (Alonso-Ramírez *et al*., 2009; Ubeda-Tomás *et al*., 2009; Lee *et al*., 2012) and down-regulate PIF4, a phytochrome-interacting factor which induces cell elongation genes (Achard and Genschik, 2009). Deficiencies in B, P or Fe, which increase the levels of DELLAs at the root MZ and induce ectopic root hairs, also reduce the primary root length (Martín-Rejano *et al*., 2011; Péret *et al*., 2011; Wild *et al*., 2016). Furthermore, it has been suggested that production of ectopic root hairs in ectopic root hair 2 *(erh2)* occurs at late stages of root development, correlated with cell expansion. However, it has also been suggested the independence of root hair initiation from cell expansion, as *erh3* acts as soon as cell fate specification (Schneider *et al*., 1997). Particularly, ERH3 is required for the stable fixation of positional signals at the cell wall (CW) for cell fate specification. Moreover, ERH3 codifies a MT-severing p60 katanin protein and has a role in CW biosynthesis (Webb *et al*., 2002). On the other hand, given that the expression of cell identity markers is altered in *erh3*, it has been suggested that MT are directly active in the specification of root cell identity, and that MT disruption in *erh3* results in the development of defective identities (Webb *et al*., 2002). In fact, in several animal systems, MT are involved in the specification of cell identity and polarity (Webb *et al*., 2002).

Shortening and radial expansion of root cortical and endodermal cells has also been reported in the *cobra, pom-1* (both cellulose deficient)*, shoebox* (GA biosynthesis-impaired), *dgl1* (GA-insensitive), TUA6/AS (α-tubulin-deficient), *erh, sabre, PLD* (phospholipase D), *wer, scm* and *jkd* mutants, in plants treated with umbelliferone (a cellulose biosynthesis inhibitor), MT-breaking drugs or 1-butanol (an inhibitor of PLD), and in plants stressed by salinity, gamma irradiation or mineral deficiency (Fe, P) (Jankay and Muller, 1976; Schiefelbein *et al*., 1997; Schneider *et al*., 1997; Bao *et al*., 2001; Ma *et al*., 2001; Scheres *et al*., 2002; Gardiner *et al*., 2003; Nagata *et al*., 2004; Komorisono *et al*., 2005; Welch *et al*., 2007; Dinneny *et al*., 2008; Pietra *et al*., 2015; Janes *et al*., 2018). Cortical cell expansion as a result of excessive DELLAs has also been described by Benfey *et al*., (1993). Interestingly, PAC induces the expression of expansin genes at the root cortex (At4g21280 (+2,002); Arex data), being *EXPANSIN 7* a specific marker of root hair cells (Ohashi *et al*., 2003; Gendre *et al*., 2019). In addition, it has been suggested that the DELLA GAI might have a role in the expansion of endodermal cells in *Arabidopsis* primary roots, and that the expansion of endodermal cells determines the elongation of whole roots (Ubeda-Tomás *et al*., 2009; Zhang *et al*., 2014). On the other hand, the GAs control the size of the root apical meristem (RAM) in *Arabidopsis* by affecting cortical cell expansion (Nelissen *et al*., 2012; Fonouni-Farde *et al*., 2019). In fact, for different accessions of *Arabidopsis*, there is a correlation between cortex cell length and the length of the root MZ (Zhang *et al*., 2014). Furthermore, when *gai-1* is over-expressed at the root endodermis (the most important tissue for GA-dependent root growth), the cessation of anisotropic cell growth expands radially the cortical cells and causes the outward protrusion of epidermal cells (Ubeda-Tomás *et al*., 2008). With this regard, it is known that the endodermis of the EZ regulates nutrient uptake (Péret *et al*., 2011; Shani *et al*., 2013; Cui, 2015), whereas the cortex participates in the root response to P deficiency (Shin *et al*., 2005).

Therefore, the shortening and widening of the root epidermal, cortical, endodermal and pericycle cells induced by excessive levels of DELLAs in seedlings of *A. thaliana* might have explained the radial expansion of the root tips observed. Interestingly, treatment with PAC also increased the root diameter in carrot (Wang *et al*., 2015). In fact, in this study, root tips became thinner under excessive GAs (Table 3), as previously reported in carrot and *Eucalyptus grandis* (Wang *et al*., 2015; Liu *et al*., 2018). A wider root diameter has also been described in the *arm, sabre, cobra, erh-1, pom-pom1* and α-tubulin under-expressing mutants, and in plants exposed to 1-butanol, umbelliferone, gamma irradiation or P deficiency (Jankay & Muller, 1976; Schneider *et al*., 1997; Bao *et al*., 2001; Ma *et al*., 2001; Gardiner *et al*., 2003; Nagata *et al*., 2004; Hermans *et al*., 2010; Pietra, 2014). Moreover, the reduction of the actin cytoskeleton induces the radial expansion of plant cells, making them shorter and wider, as the interphase MT determine the direction of plant cell elongation (Baluška *et al*., 2001; Bao *et al*., 2001). This means that the capacity of cells to elongate longitudinally depends on the orientation of the cytoskeletal MT (Dugardeyn and Van Der Straeten, 2008). Thus, a reduced expression of the α-tubulin gene in *A. thaliana* seedlings results in an abnormal expansion of the root tip (MZ and EZ), with its diameter increasing dramatically at 8 days after germination (Bao *et al*., 2001). De-polymerization of MT by oryzalin or 1-butanol also causes the swelling of the root MZ and EZ in *A. thaliana* seedlings, whereas MT stabilization by taxol expands the root EZ and DZ (Bao *et al*., 2001; Gardiner *et al*., 2003).

Therefore, a connection exists between aberrant orientation of MT and reduced cell elongation, as MT regulate the oriented deposition of cellulose microfibrills that determines the direction of cell elongation (Burk and Ye, 2002). More specifically, MT are essential for anisotropic cell expansion because they direct the insertion of cellulose synthase in the CW and guide the orientation of cellulose microfibrills to a perpendicular position with respect to the growth axis, thereby restricting radial cell expansion (Jankay and Muller 1976; Lin *et al*., 2013). Interestingly, mutations in P60 katanin protein, essential for anisotropic cell growth, cause an inappropriate feedback regulation of the *DGL1* gen for GAs biosynthesis (Komorisono *et al*., 2005).

With this respect, a link has been proposed between aberrant orientation of MT, radial cell growth and altered root hair patterning (Baluška *et al*., 2001; Bao *et al*., 2001). Thus, a low expression of the α-tubulin gene (TUA6/AS transgenic lines), mutations that inhibit MT polymerization or drugs that brake the actin MT produce aberrant microtubular structures, expand radially the root tip cells, especially at the epidermis and cortex of the MZ and EZ, and induce ectopic root hairs in 5 day-old *A. thaliana* seedlings (Bao *et al*., 2001; Collings *et al*., 2006). On the other hand, the *erh1* and *erh3* mutants, with an altered root hair patterning, exhibit disorganized MT and radially-enlarged layers of root cortex and endodermis, what suggests a connection between radial cell expansion and root hair initiation (Schneider *et al*., 1997; Bouquin *et al*., 2002; Müller and Schmidt, 2004; Pietra *et al*., 2015). Moreover, *erh2* is allelic to *pom-1*, a mutant with abnormally-expanded layers of root epidermis and cortex (Schneider *et al*., 1997; Pietra *et al*., 2015). In fact, not only cell length, but also cell width differs between trichoblasts and atrichoblasts (Löfke *et al*., 2013). Defiencies in Fe or P also induce the swelling of root cortical cells, along with ectopic root hairs (Pietra *et al*., 2015). Another clue about the link among MT, cell expansion and cell fate is illustrated by the mutants *cobra* and *sabre*, both with an abnormal cell expansion at the root tip and ectopic root hairs (Schiefelbein *et al*., 1997). The mutation of the SABRE protein, involved in MT organization, causes an abnormal cell expansion at the root cortex (Benfey *et al*., 1993), whereas the mutation of the COBRA protein, which is associated to the longitudinal CW of the rapidly-growing root EZ, entails a cellulose deficiency and causes the swelling of the root epidermis and cortex (Scheres *et al*., 2002). Interestingly, the GAs influence CW growth in mesocotyl epidermal cells (Perazza *et al*., 1998). Therefore, under the experimental conditions of the present study, excessive levels of DELLAs might have impaired the biosynthesis, organization and/or homoeostasis of MT in root tip cells of *A. thaliana* seedlings, and this, in turn, might have caused the inhibition of cell elongation and the altered patterning of GL2 and root hairs at the MZ and EZ of the root. In fact, the DELLAS destabilize the MT, giving rise to non-polar cell growth (Locascio *et al*., 2013).

In this respect, it is known that the levels of ploidy exert an important control over cell size, and that cell size and morphology are, in turn, linked to DNA content (Kondorosi *et al*., 2001). Moreover, in many tissues, cell elongation is associated to the endo-reduplication of the DNA (replication without mitosis that occurs before cell elongation, resulting in a logarithmic accumulation of genome copies in each nucleus) (Sanz *et al*., 2012). Thus, the earliest morphological signs in trichome initiation are the induction of endoreduplication and the increase in nuclear and cellular size (Perazza *et al*., 1998). With this regard, it is known that the GAs induce endo-reduplication in a dose-dependent manner and regulate cyclin gene expression. In fact, trichomes in the *spy5* mutant have two times more DNA than WT trichomes (Perazza *et al*., 1998; Kondorosi *et al*., 2001). Moreover, in GA-deficient transgenic plants, the observed root swelling, due to MT disorganization, is associated to the induction and accumulation of cyclin CYC3;1 and CYCB1;1 proteins, because the DELLAs are involved in cell cycle progression (Ubeda-Tomás *et al*., 2009; Gou *et al*., 2010; Sánchez-Calderón *et al*., 2013). In addition, the halting degree of the cell cycle is related to the GA endogenous level (Li *et al*., 2015b). Mutations in the α-tubulin gene and drugs that inhibit MT polymerization also induce multi-nucleated cells (Bao *et al*., 2001). Interestingly, the control of endoreduplication in trichomes participates in the regulation of epidermal patterning (Pietra *et al*., 2015), whereas the *RHL* genes, related to endo-reduplication, affect the fate of root epidermal cells independently from the *GL2* gene network (Guo *et al*., 2009).

Given that stress reduces root cell length and, thus, root length (Dinneny *et al*., 2008), then, the morphological alterations that were observed in the root cells under excessive levels of DELLAs in 5-day-old *A. thaliana* seedlings might be in tune with the known role of these proteins as mediators of the Stress-Induced Morphogenic Responses (SIMR) in plants, which are characterized by changes in MT metabolism, CW flexibility and cell cycle progression (Potters *et al*., 2007). Moreover, it is known that stress inhibits growth by reducing GA levels and promoting the stabilization and accumulation of DELLAs (Achard and Genschik, 2009; Alonso-Ramírez *et al*., 2009), and that the DELLAs mediate the SIMR associated to P deficiency (Jiang *et al*., 2007). From this, it might be hypothesized that an alteration of MT homoeostasis might have been implicated in the cell size changes that were observed in roots of *A. thaliana* seedlings grown under (or harbouring) excessive levels of GAs/DELLAs. While excessive levels of DELLAs might have disorganized the MT cytoskeleton, excessive levels of GAs might have stabilized it, giving rise to the changes of cell size and cell fate observed at the MZ and EZ of the root.

Results also suggest that cell fate decisions at the root epidermis might be synchronized with the cell size changes at the inner tissues of the root, such as the cortex. Thus, another reason for the appearance of extra root hairs in *A. thaliana* seedlings grown under (or harbouring) excessive levels of GAs/DELLAs might have been the reduction of the ratios of cortical/epidermal cell length (e.g. under excessive DELLAs, which decrease the cortex cell length) and of cortical/epidermal cell width (e.g. under excessive GAs, which decrease the cortex cell width), as they might give rise to epidermal cells laying over two cortical intersections instead of one, and, hence, to two-haired epidermal cells. This would imply that not only the length and width of epidermal cells, but also the length and width of cortical cells might contribute to the number of hairs produced by the root, although additional studies are needed to confirm this hypothesis.

On the other hand, results of this study also point at the cortex, endodermis and pericycle as root tissues from which the GAs/DELLAs might influence the root hair patterning, because trangenic lines over-expressing GAI at these root tissues produced ectopic root hairs and non-hairs (McCarthy-Suárez, 2021). In fact, it has been shown that blocking GAs signalling at the root endodermis induces morphological defects in the root epidermal cells (Löfke *et al*., 2013; Pietra *et al*., 2015; Janes *et al*., 2018).

#### 4.1.2. Root cell number. Connections to the root hair patterning and abundance

Excessive levels of GAs/DELLAs also altered the radial cell organization in root tips of seedlings of *A. thaliana*. While excessive levels of DELLAs frequently induced additional cells at the epidermis, cortex, endodermis and pericycle of the root, excessive levels of GAs sometimes induced extra cells at the root cortex and pericycle. With this respect, whether the cell proliferation at the root cortex-endodermis-pericycle under excessive levels of DELLAs was another reason for the observed disorganization in the root hair patterning (McCarthy-Suárez, 2021), it might be worth confirming in future experiments by using inhibitors and/or mutants of cell division.

Regarding the epidermis, interestingly, the predicted number of cells at the atrichoblast position per root radial section increased under excessive DELLAs (PAC, *gai*-1), but decreased under excessive GAs (*5X* mutant) (McCarthy-Suárez, 2021). Nevertheless, as the predicted number of cells at the trichoblast position did not change, and the percentage of ectopic root hairs was higher under excessive DELLAs as compared to excessive GAs (McCarthy-Suárez, 2021), then, the higher abundance of root hairs seen under excessive DELLAs in comparison to excessive GAs was probably due to the induction of ectopic root hairs—and, thus, to the higher number of cells at the atrichoblast position— and not because of the appearance of new trichoblast positions, given that the number of cells at the trichoblast position remained unchanged (McCarthy-Suárez, 2021).

In fact, *Arabidopsis* increases root hair density in the radial dimension by increasing the number of epidermal cells that differentiate into root hair cells (Janes *et al*., 2018). An increased number of epidermal cells in the radial domain of the root has also been described under P deficiency and in *tip1* mutants (Ma *et al*., 2001; Grierson and Schiefelbein, 2002; Müller and Schmidt, 2004). However, under P deficiency, the extra epidermal cells at the trichoblast position (up to 12) do not increase the abundance of root hairs in the radial axis, as ectopic non-hairs also appear (Ma *et al*., 2001; Janes *et al*., 2018). Interestingly, a distorted radial patterning of root cells has also been described in mutants of WRKY75, a negative regulator of root hair formation (Rishmawi *et al*., 2014).

Radial proliferation of the root cortex cells has also been reported under stress (e.g. P deficiency) as well as in *tip1, erh3* and *jkd* mutants, all with an altered root hair patterning (Ma *et al*., 2001; Müller and Schmidt, 2004; Hassan *et al*., 2010; Cui, 2015; Janes *et al*., 2018). Periclinal cell divisions (extra layers) of the root cortex have equally been described in mutants of JKD, which acts from the root cortex to specify the patterning of epidermal cell types (Welch *et al*., 2007; Lyer-Pascuzzi and Benfey, 2008; Hassan *et al*., 2010). Interestingly, the GAs restrict the production of extra cortex cell layers in *Medicago truncatula* roots, thereby generating thinner roots (Fonouni-Farde *et al*., 2019). In contrast, PAC treatments, or mutations in components of GA signalling, increase the number of layers of root cortex cells, that is, they induce a premature middle cortex (MC) (Paquette and Benfey, 2005; Cui & Benfey, 2009). Moreover, the GAs suppress the MC formation that is proper of the root responses to stress, whereas the DELLAs promote it (Cui and Benfey, 2009; Fonouni-Farde *et al*., 2019). Thus, the formation of a MC, due to random and periclinal cell divisions at the root endodermis, and that later on will acquire identity of root cortex, has been described under P deficiency (Cui and Benfey, 2009; Janes *et al*., 2018). Although the production of a MC has also been reported in roots of 3-day-old WT *A. thaliana* seedlings, the presence of a premature MC in the *spy* mutant, with high levels of GAs, suggests that imbalances in GAs/DELLAs homoeostasis, which can be triggered by stress, might bring about the formation of a MC (Cui and Benfey, 2009; Cui, 2015). Maybe this was the reason, in this study, for the presence of a MC in roots of *A. thaliana* seedlings grown under (or harbouring) excessive levels of GAs (1 μM GA_4_ and *QD*) (Figs. 5 and 6).

Because the PAC-inducible MC phenotype is also present in the *scr* (scarecrow) and HDA mutants, as well as in trichostatin A (TSA)-treated plants, all producing ectopic root hairs (Cui and Benfey, 2009), then, a possible link between MC formation (or ectopic cell proliferation at the cortex/endodermis) and alteration of the root hair patterning might be established. Whether this contributed to the disorganisation of the root hair patterning observed under excessive levels of DELLAs, where a cell proliferation was equally observed at the cortex/endodermis/pericycle of the root, it is not known, but might be worth studying in future experiments by using cell division inhibitors and/or mutants. In fact, the alteration of the root hair patterning in the *SCR::gai-1:GR* mutant after growth in DEXA (McCarthy-Suárez, 2021) was accompanied by random and periclinal cell divisions at the root endodermis (Fig. 3). Moreover, because the root MZ and EZ constitute cell fate-decision zones in *Arabidopsis*, then, any changes in cell division at tissues placed underneath the epidermis of the MZ/EZ might bring about changes in epidermal cell fate. In fact, the proliferation of cortex cells is known to influence the root epidermal patterning (Löfke *et al*., 2013; Pietra *et al*., 2015; Janes *et al*., 2018). Interestingly, histone deacetylation, which affects the root hair patterning, has a role in the proliferation of root cortex cells (Xu *et al*., 2005; Li *et al*., 2015a). Increases in the number of root endodermal cells have also been reported in *erh* mutants, during P deficiency, and in *rhizobium-infected* plants (Müller and Schmidt, 2004; Ma *et al*., 2001; Janes *et al*., 2018). The schizorriza (*scz*) mutant, in turn, has defects in the root radial patterning, with extra periclinal cell divisions that result in multiple layers of ground tissue (cortex and endodermis) (Mylona *et al*., 2002). Thus, the results of this study suggest that the alterations in the root hair patterning of *A. thaliana* seedlings grown under excessive levels of GAs/DELLAs might also have been related to changes in the number of the cortical/endodermal/pericycle cells of the root.

Other possible cause, in this study, for the appearance of ectopic root hairs might have been the anticlinal, diagonal or asymmetric cell divisions (T-clones) frequently observed under excessive levels of GAs/DELLAs at the root epidermis, as they gave rise to changes in the *GL2pro::GUS* patterning and the size of daughter cells (Fig. 3). These T-clones, in turn, might have been linked to alterations in the MT cytoskeleton, as MT are required for the correct positioning of cell division planes (Scheres and Benfey, 1999; Bao *et al*., 2001; Rodriguez-Serrano *et al*., 2014). Moreover, a reduced expression of the α-tubulin gene impairs cell division and results in defects of tissue organization at the root tip (Bao *et al*., 2001). Also, the regulation of asymmetric cell divisions in plants is necessary for the generation of cell diversity and patterns (Pernas *et al*., 2010). In fact, root hair cells are shorter than root non-hair cells, so that when an asymmetric cell division takes place at the epidermis of the MZ, the larger cell becomes the root non-hair cell (Salazar-Henao *et al*., 2016). With this respect, it is known that the GAs induce cell proliferation at the root MZ and promote the division of epidermal cells (Ubeda-Tomás *et al*., 2009; Lee *et al*., 2012). Furthermore, the DELLAs inhibit root cell division in the longitudinal dimension when mediating the SIMR associated to P deficiency (Jiang *et al*., 2007; Péret *et al*., 2011). Interestingly, in a dwarf GA-deficient mutant, the MT exhibit a oblique orientation (Bouquin *et al*., 2002). Also, in the *erh3* mutants, which act in the same route as *cpc* and *rhd6* (root hair defective 6), but independently from WER, the CW are dis-aligned, diagonally orientated, and malformed, that is, the positioning of the cell plates and the CW is abnormal, what indicates that ERH3 participates in orienting the cell plates during cytokinesis (Webb *et al*., 2002). The cortex-associated SABRE protein is also involved in the orientation of cell division planes (Pietra, 2014; Pietra *et al*., 2013, 2015). In addition, GL2 is involved in the production of T-clones, whereas WER regulates cell proliferation, what suggests that genes that regulate cell specification also regulate cell division planes (Lee & Schiefelbein, 1999; Scheres *et al*., 2002).

Thus, these results suggest that changes in cell number induced by excessive levels of GAs/DELLAs at the epidermis, cortex, endodermis and pericycle of the root tip in *A. thaliana* seedlings might influence the root hair patterning. Moreover, changes in cell number at tissues placed underneath the root epidermis might bring changes of cell fate at the root epidermal cells.

In addition, results of this study suggest that excessive levels of DELLAs in roots of *A. thaliana* seedlings might impair the biosynthesis and/or the assembly of MT, as judged by the swelling of the root tip cells (Figs. 1–4), the presence of multi-nucleated cells at the MZ (Fig. 6), as well as by the occurrence of ectopic root hairs, branched root hairs, and cells with multiple root hairs (McCarthy-Suárez, 2021). In fact, a reduced expression of the α-tubulin gene results in the disassembly and aberrant reorganization of MT (Bao *et al*., 2001). This means that the alteration of the root hair patterning in *A. thaliana* seedlings by excessive levels of DELLAs might have been correlated to their inhibitory effect on MT organisation. In fact, MT are essential to stablish root cell identity in *Arabidopsis* (Webb *et al*., 2002).

### 4.2. The GAs/DELLAs might regulate the root architecture in *A. thaliana* seedlings

Excessive levels of DELLAs also promoted the outburst of LR near the root tip in *A. thaliana* seedlings, whereas excessive levels of GAs inhibited it. This means that any alteration in the levels of GAs/DELLAs might affect not only the root hair patterning, morphology and abundance, but also the root architecture. Thus, in seedlings of *A. thaliana*, physiologically-controlled levels of GAs/DELLAs might have a function in establishing a correct patterning and morphology of root hairs as well as in organising a proper root structure. In fact, supra-physiological levels of DELLAs mediate the root architecture changes that are associated to abiotic stress in plants (i.e., root elongation inhibition, root radial expansion, MC formation, pericycle cell proliferation and LR induction) (Yih and Clark, 1965; Jiang *et al*., 2007; Gou *et al*., 2010; Martín-Rejano *et al*., 2011; Péret *et al*., 2011; Cui, 2015; Wild *et al*., 2016). For example, soil deficiencies of P, B, Fe or NO_3_^-^ stimulate LR production, as nutrient concentration regulates LR production (Yih and Clark, 1965; Hermans *et al*., 2010; Zhang *et al*., 2014). Early production of LR has also been reported in the *erh1, jkd* and *arm* mutants, equally with a shortened primary root (Schneider *et al*., 1997; Welch *et al*., 2007). Interestingly, LR formation, which initiates at the pericycle, is correlated to an alteration in actin and tubulin expression (Pasternak *et al*., 2005; Péret *et al*., 2011; Sánchez-Calderón *et al*., 2013). Therefore, promoting LR outburst by increasing the local levels of DELLAs in roots might constitute a mechanism used by plants to increase the specific area of the root per mass unit, in a similar way as branched root hairs do.

In conclusion, as it was previously reported for other hormones, the results of this study, and of a previous study (McCarthy-Suárez, 202?), point to a possible role for the GAs/DELLAs in mediating the changes in the distribution, shape and frecuency of root hairs, as well as in the root configuration, that take place in plants under stress situations. Moreover, the auxins, ET, ABA, BRs and SLs mediate these changes without altering the quantitative expression of *WER* and *GL2* (Schiefelbein, 2003; Yang *et al*., 2007; Martín-Rejano *et al*., 2011). This implies that, by regulating the elongation and/or division of root cells, as well as the production of LR, the GAs/DELLAs might be potential mediators of the changes in the root hair patterning, morphology and abundance, and of the changes in the root architecture, occurring in plants under environmental stress conditions.

## Acknowledgements

Special thanks to Dr. Miguel Ángel Blázquez and Dr. David Alabadí for supporting the writing of this paper. This study was performed at the Blázquez-Alabadí Laboratory (Hormone Signalling and Plant Plasticity Group, Instituto de Biología Celular y Molecular de Plantas (IBMCP)-CSIC-UPV, Valencia, Spain). Thanks also to Mrs. Marisol Gascón Irún for confocal microscope technical support. I. McCarthy Suárez also acknowledges a JAE-Doc postdoctoral fellowship (2008-2011), CSIC, Valencia, Spain. This paper is dedicated to the loving memory of Dr. Francisco Culiáñez-Macià, Principal Investigator at the IBMCP-UPV-CSIC, Valencia, Spain.

## References

Achard P and Genschik P (2009) Releasing the brakes of plant growth: how GAs shutdown DELLA proteins. Journal of Experimental Botany 60: 1085–1092.

Alonso-Ramírez A, Rodríguez D, Reyes D, Jiménez JA, Nicolás G, López-Climent M, Gómez- Cárdenas A and Nicolás C (2009) Evidence for a role of gibberellins in salicylic acid-modulated early plant responses to abiotic stress in *Arabidopsis* seeds. Plant Physiology 150: 1335–1344.

Bahin E, Bailly C, Sotta B, Kranner I, Corbineau F and Leymarie J (2011) Crosstalk between reactive oxygen species and hormonal signalling pathways regulates grain dormancy in barley. Plant, Cell and Environment 34: 980–993.

Baluška F, Jasik J, Edelmann HG, Salajová T and Volkmann D (2001) Latrunculin B-induced plant dwarfism: plant cell elongation is F-actin dependent. Developmental Biology 231: 113–124.

Band LR, Ubeda-Tomás S, Dyson RJ, Middleton AM, Hodgman C, Owen MR, Jensen OE, Bennett MJ, King JR (2012) Growth-induced hormone dilution can explain the dynamics of plant root cell elongation. Proceedings of the National Academy of Sciences (PNAS) 109: 7577–7582.

Bao Y, Kost B and Chua NH (2001) Reduced expression of α-tubulin genes in *Arabidopsis thaliana* specifically affects root growth and morphology, root hair development and root gravitropism. The Plant Journal 28: 145–157.

Benfey PN, Linstead PJ, Roberts K, Schiefelbein JW, Hauser MT and Aesbacher RA (1993) Root development in *Arabidopsis*: Four mutants with dramatically altered root morphogenesis. Development 119: 57–70.

Bouquin T, Mattson O, Naested H, Foster R and Mundy J (2002) The *Arabidopsis lue1* mutant defines a katanin p60 ortholog involved in hormonal control of microtubule orientation during cell growth. Journal of Cell Science 116: 791–801.

Burk DH and Ye ZH (2002) Alteration of oriented deposition of cellulose microfibrills by mutation of a katanin-like microtubule severing protein. The Plant Cell 14: 2145–2160.

Cao XF, Linstead P, Berger F, Kieber J and Dolan L (1999) Differential ethylene sensitivity of epidermal cells is involved in the establishment of cell pattern in the *Arabidopsis* root. Physiologiae Plantarum 106: 311–317.

Chen CY, Wu K and Schmidt W (2015) The histone deacetilase HDA19 controls root cell elongation and modulates a subset of phosphate starvation responses in *Arabidopsis*. Scientific reports 5: 15708. DOI: 10.1038/srep15708.

Chien JC and Sussex IM (1996) Differential regulation of trichome formation on the adaxial and abaxial surfaces by gibberellins and photoperiod in *Arabidopsis thaliana* (L.) heynh. Plant Physiology 111: 1321–1328.

Collings DA, Lill AW, Himmelspach R and Wasteneys GO (2006) Hypersensitivity to cytoskeletal antagonists demonstrates microtubule-microfilament cross-talk in the control of root elongation in *Arabidopsis thaliana*. New Phytologist 170: 275–290.

Cui H (2015) Cortex proliferation in the root is a protective mechanism against abiotic stress. Plant Signalling and behaviour 10:5, e1011949.

Cui H and Benfey P (2009) Interplay between scarecrow, GA and like heterochromatin protein 1 in ground tissue patterning in the *Arabidopsis* root. The Plant Journal 58: 1016–1027.

Dinneny JR, Long TA, Wang JY, Jung JW, Mace D, Pointer S, Barron C, Brady SM, Schiefelbein J and Benfey PN (2008) Cell identity mediates the responses of *Arabidopsis* roots to abiotic stress. Science 320: 942–945.

Dugardeyn J and Van der Straeten D (2008) Ethylene: Fine tuning plant growth and development by stimulation and inhibition of elongation. Plant Science 175: 59–70.

Fonouni-Farde C, Miassod A, Laffont C, Morin H, Bendahmane A, Diet A and Frugier F (2019) Gibberellins negatively regulate the development of *Medicago truncatula* root system. Scientific Reports 9: 2335.

Gardiner J, Collings DA, Harper JDI and Marc J (2003) The effects of the phospholipase D-antagonist 1- butanol on seedlings development and microtubule organisation in *Arabidopsis*. Plant Cell Physiology 44: 687–696.

Gendre D, Baral A, Dang X, Esnay N, Boutté Y, Stanisla S, Vain T, Claverol S, Gustavsson A, Lin D, Grebe M and Bhalerao R (2019) Rho-of-plant activated root hair formation requires *Arabidopsis* YIP4alb gene function. Development 146, dev 168559, DOI: 10.1242/dev.168559.

Gou J, Strauss SH, Tsai CJ, Fang K, Chen Y, Jiang X and Busov VB (2010) Gibberellins regulate lateral root formation in *Populus* through interactions with auxin and other hormones. The Plant Cell 22: 623–639.

Grierson C and Schiefelbein J (2002) Root hairs. The Arabidopsis book 2–22. C.R. Somerville, E.M. Meyerowitz, Eds. Publisher: American Society of Plant Biologists. Rockville, MD.

Guo K, Kong WW and Yang ZM (2009) Carbon monoxide promotes root hair development in tomato. Plant, Cell & Environment 32: 1033–1045.

Hassan H, Scheres B and Blilou I (2010) Jackdaw controls epidermal patterning in the *Arabidopsis* root meristem through a non-cell autonomous mechanism. Development 137: 1523–1529.

Hermans C, Porco S, Verbruggen N and Bush DR (2010) Chitinase-like protein CTL-1 plays a role in altering root system architecture in response to multiple environmental conditions. Plant Physiology 152: 904–917.

Janes G, von Wangenheim D, Cowling S, Kerr I, Band L, French AP and Bishop A (2018) Cellular patterning of *Arabidopsis* roots under low phosphate conditions. Frontiers in Plant Science 9: 735.

Jankay P and Muller WH (1976) The relationship among umbelliferone, growth, and peroxidase levels in cucumber roots. American Journal of Botany 63: 126–132.

Jiang C, Gao X, Liao L, Harberd NP and Fu X (2007) Phosphate starvation root architecture and anthocyanin accumulation responses are modulated by the gibberellin-DELLA signalling pathway in *Arabidopsis*. Plant Physiology 145: 1460–1470.

Kappusamy KT, Chen AY and Nemhauser JL (2009) Steroids are required for epidermal cell fate establishment in *Arabidopsis* roots. Proceedings of the National Academy of Sciences (PNAS) 106: 8073–8076.

Komorisono M, Ueguchi-Tanaka M, Aichi I, Hasegawa Y, Ashikari M, Kitano H, Matsuoka M and Sazuka T (2005) Analysis of the rice mutant *dwarf* and *gladius leaf1* aberrant katanin-mediated microtubule organization causes up-regulation of gibberellin biosynthetic genes independently of gibberellin signalling. Plant Physiology 138: 1982–1993.

Kondorosi E, Roudiera F and Gendreau E (2001) Plant cell size control: growing by ploidy? Current Opinion in Plant Biology 3: 488–492.

Lee MM and Schiefelbein J (1999) WEREWOLF, a MYB-related protein in *Arabidopsis*, is a position-dependent regulator of epidermal cell patterning. Cell 99: 473–483.

Lee LY, Hou X, Fang L, Kumar PP and Yu H (2012) Stunted mediates the control of cell proliferation by GA in *Arabidopsis*. Development 139: 1568–1576.

Li DX, Chen WQ, Xu ZH and Bai SN (2015)a Histone deacetilase 6-defective mutants show increased expression and acetylation of enhancer of tryptychon and caprice1 and glabra2 with small but significant effects on root epidermis cellular pattern. Plant Physiology 168: 1448–1458.

Li J, Zhao Y, Chu H, Wang L, Fu Y, Liu P, Upadhyaya N, Chen C, Mou T, Feng Y, Kumar P and Xu J (2015)b Shoebox modulates root meristem size in rice through dose-dependent effects of gibberellins on cell organisation and proliferation. PLOS GENETICS DOI: 10.1371/journal.pgen.1005464.

Lin D, Cao L, Zhou Z, Zhu L, Ehrhardt D, Yang Z and Fu Y (2013) Rho GTPase signalling activates microtubule severing to promote microtubule ordering in *Arabidopsis*. Current Biology 23: 290–297.

Liu QY, Guo GS, Qiu ZF, Li XD, Zeng BS and Fan CJ (2018) Exogenous GA3 application altered morphology, anatomic and transcriptional regulatory networks of hormones in *Eucalyptus grandis*. Protoplasma 255: 1107–1119.

Locascio A, Blázquez MA and Alabadí D (2013) Dynamic regulation of cortical microtubule organization through prefoldin-DELLA interaction. Current Biology 23: 804–809.

Löfke C, Dünster K and Kleine-Vehn J (2013) Epidermal patterning genes impose non-cell autonomous cell size determination and have additional roles in root meristem size control. Journal of Integrative Plant Biology 55: 864–875.

Lombardo MC, Graziano M, Polacco JC and Lamattina L (2006) Nitric oxide functions as a positive regulator of root hair development. Plant Signalling and Behaviour 1: 28–33.

Lyer-Pascuzzi A and Benfey PN (2008) Transcriptional networks in root cell fate specification. Biochimica et Biophysica Acta 1789: 315–325.

Ma Z, Bielenberg GD, Brown KM and Lynch JP (2001) Regulation of root hair density of phosphorus availability in *Arabidopsis thaliana*. Plant Cell & Environment 24: 459–467.

Martín-Rejano EM, Camacho-Cristóbal JJ, Herrera-Rodríguez MB, Rexach J and Navarro-Gochicoa MT (2011) Auxin and ethylene are involved in the responses of root system architecture to low boron supply in *Arabidopsis* seedlings. Physiologia Plantarum 142: 170–178.

McCarthy-Suárez I (2021) Supra-physiological levels of Gibberellins/DELLAs alter the patterning, morphology and abundance of root hairs in root tips of *A. thaliana* seedlings. BioRxiv/2021/452505.

Mylona P, Linstead P, Martienssen R and Dolan L (2002) Schizorriza controls an assymetric cell division and restricts epidermal identity in the *Arabidopsis* root. Development 129: 4327–4334.

Müller M and Schmidt W (2004) Environmentally induced plasticity of root hair development in *Arabidopsis*. Plant Physiology 134: 409–419.

Nagata T, Todoriki S and Kikuchi (2004) Radial extension of root cells and elongation of root hairs of *Arabidopsis thaliana* induced by massive doses of gamma irradiation. Plant Cell Physiology 45: 1557–1565.

Nelissen H, Rymen B, Jikumaru Y, Demuynk K, Van Lijsebettens M, Kamiya Y, Inzé D and Beemster GTS (2012) A local maximum in gibberellin levels regulates maize leaf growth by spatial control of cell division. Current Biology 22: 1183–1187.

Niu Y, Jin C, Jin G, Zhou Q, Lin X, Tang C and Zhang Y (2011) Auxin modulates the enhanced development of root hairs in *Arabidopsis thaliana* (L.) Heyhn. under elevated CO(2). Plant, Cell and Environment 34: 1304–1317.

Ohashi Y, Oka A, Rodrígues-Pousada R, Possenti M, Ruberti I, Morelli G and Aoyama T (2003) Modulation of phospholipid signalling by GLABRA2 in root hair pattern formation. Science 300: 1427–1430.

Paquette AJ and Benfey PN (2005) Maturation of the ground tissue of the root is regulated by gibberellin and scarecrow and requires short-root. Plant Physiology 138: 636–640.

Pasternak T, Potters G, Caubergs R and Jansen MAK (2005) Complementary interactions between oxidative stress and auxins control plant growth responses at plant, organ and cellular level. Journal of Experimental Botany 56: 1991–2001.

Perazza D, Vachon G and Herzog M (1998) Gibberellins promote trichome formation by up-regulating *GLABROUS1* in *Arabidopsis*. Plant Physiology 117: 375–383.

Péret B, Clément M, Nussaume L and Desnos T (2011) Root developmental adaptation to phosphate starvation: Better safe than sorry. Trends in Plant Science 16: 442–450.

Pernas M, Ryan E and Dolan L (2010) Schizorrhyza controls tissue system complexity in plants. Current Biology 20: 812–823.

Pietra S (2014) Characterization of new players in planar polarity establishment in *Arabidopsis*. Doctoral thesis Umea Plant Science Centre Fysiologisk Botanik.

Pietra S, Gustavsson A, Kiefer C, Kalmbach L, Hörstedt P, Ikeda Y, Stepanova AN, Alonso JM and Grebe M (2013) *Arabidopsis* SABRE and CLASP interact to stabilize cell division plane orientation and planar polarity. Nature Communications 4: 2779.

Pietra S, Lang P and Grebe M (2015) *SABRE* is required for stabilization of root hair patterning in *Arabidopsis thaliana*. Physiologiae Plantarum 153: 440–453.

Potters G, Pasternak TP, Guisez Y, Palme KJ and Jansen MAK (2007) Stress-induced morphogenic responses: Growing out of trouble? Trends in Plant Science 12: 98–105.

Rishmawi L, Pesch M, Juengst C, Schauss AC, Schrader A and Hülskamp M (2014) Non-cell autonomous regulation of root hair patterning genes by WRKY75 in *Arabidopsis*. Plant Physiology 165: 186–195.

Rodríguez-Serrano M, Pazmiño DM, Sparkes I, Rochetti A, Hawes C, Romero-Puertas and Sandalio LM (2014) 2,4-dichlorophenoxyacetic acid promotes S-nitrosylation and oxidation of actin affecting cytoskeleton and peroxisomal dinamics. Journal of Experimental Botany 65: 4783–4793.

Salazar-Henao JE, Vélez-Bermúdez IC and Schmidt W (2016) The regulation and plasticity of root hair patterning and morphogenesis. Development 143: 1848–1858.

Sánchez-Calderón L, Ibarra-Cortés ME and Zepeda-Jazo I (2013) Root development and abiotic stress adaptation. Abiotic stress-plant responses and applications in agriculture. Chapter 5 135–168. Intech Open Science.

Sanz L, Murray JAH and Dewitte W (2012) To divide and to rule: Regulating cell division in roots during post-embryonic growth. Progress in Botany 73: 57–80.

Scheres B and Benfey PN (1999) Asymmetric cell division in plants. Annual Review of Plant Physiology and Plant Molecular Biology 50: 505–537.

Scheres B, Benfey P and Dolan L (2002) Root development. The Arabidopsis book. C.R. Somerville, E.M. Meyerowitz, Eds. Publisher: American Society of Plant Biologists, Rockville, MD.

Schiefelbein J (2003) Cell fate specification in the epidermis: A common patterning mechanism in the root and the shoot. Current opinion in Plant Biology 6: 74–78.

Schiefelbein J, Masucci JD and Wang H (1997) Building a root: The control of patterning and morphogenesis during root development. The Plant Cell 9: 1089–1098.

Schmidt W, Tittel J and Schikora A (2000) Role of hormones in the induction of iron deficiency responses in *Arabidopsis* roots. Plant Physiology 122: 1109–1118.

Schneider K, Wells B, Dolan L and Roberts K (1997) Structural and genetic analysis of epidermal cell differentiation in *Arabidopsis* primary roots. Development 124: 1789–1798.

Shani E, Weinstain R, Zhang Y, Castillejo C, Kaiserli E, Chory J, Tsien RY and Estelle M (2013) Gibberellins accumulate in the elongating endodermal cells of *Arabidopsis* root. Proceedings of the National Academy of Sciences (PNAS) USA 110: 4834–4839.

Shin LJ, Huang HE, Chang H, Lin YN, Feng TY and Ger MJ (2011) Ectopic ferredoxin I protein promotes root hair growth through induction of reactive oxygen species in *Arabidopsis thaliana*. Journal of Plant Physiology 168: 434–440.

Traw MB and Bergelson J (2003) Interactive effects of jasmonic acid, salicylic acid, and gibberellin on induction of trichomes in *Arabidopsis*. Plant Physiology 133: 1367–1375.

Ubeda-Tomás S, Swarup R, Coates J, Swarup K, LaPlace L, Beemster GTS, Hedden P, Bhalerao R and Bennett MJ (2008) Root growth in *Arabidopsis* requires gibberellin/DELLA signalling in the endodermis. Nature Cell Biology 10: 625–628.

Ubeda-Tomás S, Federici F, Casimiro I, Beemster GT, Bhalerao R, Swarup R, Doerner P, Hasselhoff J and Bennett MJ (2009) Giberellin signalling in the endodermis controls *Arabidopsis* root meristem size. Current Biology 19: 1194–1199.

Van Hengel AJ, Barber C and Roberts K (2004) The expression patterns of arabinogalactan-protein AtAGP30 and GLABRA2 reveal a role for abscisic acid in the early stages of root epidermal patterning. The Plant Journal 39: 70–83.

Wang GL, Que F, Xu ZS, Wang F and Xiong AS (2015) Exogenous gibberellin altered morphology, anatomic and transcriptional regulatory networks of hormones in carrot root and shoot. BMC Plant Biology 15:290.

Webb M, Jouannic S, Foreman J, Linstead P and Dolan L (2002) Cell specification in the *Arabidopsis* root epidermis requires the activity of ECTOPIC ROOT HAIR 3-a katanin P60 protein. Development 129: 123–131.

Welch D, Hassan H, Blilou I, Immink R, Heidstra R and Scheres B (2007) *Arabidopsis* JACKDAW and MAGPIE zinc finger proteins delimit asymmetric cell division and stabilize tissue boundaries by restricting short-root action. Genes and Development 21: 2196–2204.

Wild M, Davière JM, Regnault T, Dubeaux G, Vert G and Achard P (2016) Tissue-specific regulation of gibberellin signaling fine-tunes *Arabidopsis* iron-deficiency responses. Developmental Cell 37: 190–200.

Xu CR, Liu C, Wang YL, Li LC, Chen WQ, Xu ZN and Bai SN (2005) Histone deacetylation affects expression of cellular patterning genes in the *Arabidopsis* root epidermis. Proceedings of the National Academy of Sciences (PNAS) 20: 14469–14474.

Yang T, Savage N and Schmidt W (2007) Plasticity of root epidermal cell fate in response to nutrient starvation. 18th International Conference on Arabidopsis Research.

Yih RY and Clark HE (1965) Carbohydrate and protein content to boron-deficient tomato root tips in relation to anatomy and growth. Plant Physiology 40: 312–315.

Zhang C, Bousquet A and Harris JM (2014) Abscisic acid and lateral root organ defective/numerous infections and polyphenolics modulate root elongation via reactive oxygen species in *Medicago truncatula*. Plant Physiology 166: 644–658.

